# Genome-wide mapping of the *Galleria mellonella* larvae transcription start sites during fungal infection and treatment

**DOI:** 10.1101/2025.01.20.627872

**Authors:** Imad Abugessaisa, Mickey Konings, Ri-ichiroh Manabe, Michihira Tagami, Jessica Severin, Akira Hasegawa, Tsugumi Kawashima, Hiroko Kinoshita, Shohei Noma, Chitose Takahashi, Annelies Verbon, Yasushi Okazaki, Wendy W.J. van de Sande, Takeya Kasukawa

## Abstract

Using Low Quantity single strand CAGE (LQ-ssCAGE), we mapped the transcription start sites (TSS). We annotated the 5’ end of the invertebrate *Galleria mellonella,* an upcoming and booming experimental model in infectious disease and immunology research. However, the current genome annotation of this model organism lacks annotation of the 5’ end and TSS information. *G. mellonella* larva was infected with the fungal pathogen *Madurella mycetomatis to map TSS under healthy and infection conditions*. Larvae were first treated with itraconazole or ravuconazole, and then RNA-seq and LQ-ssCAGE libraries were prepared and sequenced 4, 30, and 52 hours following infection. The LQ-ssCAGE data was processed to identify CAGE transcription start site (CTSS), uni-, and bi-directional clusters. LQ-ssCAGE enabled us to precisely identify 39,410 TSS and 249 active enhancers; we assigned genomic features to the resulting TSSs and enhancers. The majority of the TSS peaks are annotated as promoter regions, while the enhancers were annotated as intergenic and genic. Furthermore, we confirmed the quality of TSS calling by promoter shapes and GC bias. Furthermore, we identified a set of super-enhancers and predicted de-novo motifs. The raw and processed data was deposited to NCBI GEO GSE282923. CTSS, TSS peaks, and enhancers coordinated are available through the ZENBU Genome browser. In this study, we reported the first atlas of TSS and active enhancers of *G. mellonella*.

## Background

The greater wax moth *Galleria mellonella* (Linnaeus, 1758) is an invertebrate that belongs to the Pyralidae family. Recently, *G. mellonella* has become an attractive disease model for biological research (**Supplementary Figure 1**). Using the larvae of *G. mellonella*, several in vivo infection studies were performed, including disease modeling, host-pathogen interaction, the innate immune response, and testing the efficacy of novel drugs via susceptibility test[1]. As an infection model[2], the model enabled the study of infectious disease phenotypes that are difficult to study and characterize in human patients[2]. Using *G. mellonella* as a disease model in biological research has several benefits. First, it is cheap and easy to host, has a short life cycle with many offspring, and it doesn’t require ethical clearance. As a research model, *G. mellonella* reduces the use of vertebrates in biomedical research, which is much more costly.

The first genome of *G. mellonella* was sequenced in 2017 and contained 1,937 contigs (Genome assembly ASM258982v1), this was followed by several initiatives to enhance the genome assembly and provided gene annotation utilizing the latest development in long-read sequencing. The current gene set of *G. mellonella* includes 89.9% protein-coding genes, 5.6% ncRNA, 3.2% small RNA, and 1.3% pseudogenes[3]. The availability of a high-quality genome assembly will enable studies to understand the structure and function of the genome model[4]. Genome annotation can either be obtained from public databases or via computational analysis and prediction.

Together with the availability of reference genomes, gene and feature annotations are provided by Ensembl[5] and NCBI[6]. Novel biological insights, like understanding transcriptional regulation and RNAs production, can be obtained by utilizing these resources. The instructions for gene regulation determine when, where, and at which level a gene or set of genes are transcribed in a specific cell; these instructions are encoded in cis-regulatory elements (CREs). Promoters and enhancers are CREs and contain transcription factor (TF) binding sites (TFBS). The gene expression levels in cell types are determined by the TFs together with different transcriptional regulators[7]. In general, CREs can be divided into two categories: constitutively open CREs and CREs accessible in a cell-type-specific manner[8]. The transcription start site (TSS) is the specific genomic region in which transcription is initiated. TSSs are dual-regulated by TFs and promoters[9] and the genomic region of the TSS in the genome makes it the most suitable unit to integrate various types of data related to transcriptional events[9]. Mapping and characterization of TSSs are required to understand transcription regulation and to understand how TFs, chromatin proteins, and TFBSs interact with promoter regions to regulate transcription[10].

Methods for mapping TSSs began with primer extension studies[11] followed by rapid amplification of cDNA ends for example RACE, which is a technique used to clone full-length transcript and transcript variants[12, 13], as well as the expressed sequence tag (EST) approach for cap-trapped 5′ end expressed sequence tag sequencing[14]. The most recent methods to map the 5′ end cap of the transcript, cap analysis of gene expression (CAGE)[15] and its subsequent developments, no-amplification non-tagging CAGE (nAnT-iCAGE[16]), nano-cap analysis of gene expression (nanoCAGE)[17], and Low Quantity Single Strand CAGE (LQ-ssCAGE)[18], enabled the mapping of TSSs and identification of promoters and their activities. In addition to the TSS peaks, CAGE data allows for the prediction of active enhancers, or bi-directional clusters of CAGE-TSS (CTSS). With the proceeding information in mind, the main analysis of CAGE data involves the clustering of CAGE tags, such analysis considers identifying a group of CTSS clusters corresponding to the expression activity of a single TSS and related enhancers[19].

The lack of available TSS information for *G. mellonella* motivated us to map TSS information, and identify *G. mellonella* cis-regulatory elements (CREs). (promoters and active enhancers).

To perform genome-wide mapping and annotation of TSSs in *G. mellonella* we used RNA isolated from *G. mellonella* larvae then prepared and sequenced LQ-ssCAGE libraries (*n*=72). We prepared and sequenced RNA-seq from similar samples to validate the quality of the LQ-ssCAGE mapping. LQ-ssCAGE identifies the TSSs of the 5’ end capped RNA, and at the same time, measures the expression level of the transcript[18]. *G. mellonella* is an effective model for analyzing disease burden and response to treatment; we infected *G. mellonella* larvae with the fungal pathogen *M. mycetomatis* (NCBI:txid100816). *M. mycetomatis* belongs to the order of the Sordariales in the Sordariomycetes family and is the most common causative agent of Eumycetoma. Eumycetoma is a Neglected Tropical Disease affecting the subcutaneous tissue[20]. To cure eumycetoma, patients are usually treated with a combination of antifungal treatment and surgery[21]. Currently, the antifungal agent itraconazole is the drug of choice, along with ravuconazole which was a drug recently assessed in the first clinical trial for eumycetoma[22]. Therefore, to mimic a natural infection and treatment, we divided the *M. mycetomatis-*infected larvae into three groups. One control group was treated with solvent only, one group was treated with itraconazole, and one group with ravuconazole. LQ-ssCAGE (*n*=72) and RNA-seq (*n*=72) libraries were deeply sequenced using next-generation sequencing platforms, which resulted in a large 5’ end, promoter-rich dataset for *G. mellonella*. The LQ-ssCAGE data enables us to identify CTSSs; TSSs, characterized by uni- directional clusters; and enhancers, characterized by bi-directional clusters. We analyzed the TSS shapes, either sharp or broad, and confirmed the known quality of promoters, including the GC content bias in promoter regions. We identified interacting TSSs and enhancers as TSSs and enhancers that are both in proximity to each other, and highly correlate in their expression[23]. Our analysis aimed to confirm the general features of the enhancers, to use LQ-ssCAGE to analyze differential TSS usage, and using these TSS peaks, we performed de-novo motif analysis and analyzed the variation of the expression of the predicted TFs between healthy and infected samples.

We provide a genomic view of *G. mellonella* new annotation via the ZENBU genome browser[24]. This enables the research community to browse the genomic location of the TSS peaks and enhancers visually. Also, users will be able to upload any gene expression data generated from *G. mellonella* to the ZENBU genome browser moving forward.

Our genome-wide mapping and annotation of *G. mellonella* larvae TSS peaks will contribute to the gene annotation of *G. mellonella* and encourage the continued usage of the model in biomedical research.

## RESULTS

### CAGE tag 5’ end quality and coverage

We extracted total RNA from 72 time-course samples under different conditions (healthy, infected, healthy treated, infected treated), with three biological replicates for each condition (**Figure 1A**). The extracted total RNA was used for the preparation of two types of sequencing libraries RNA-seq and LQ-ssCAGE (**METHODS**). For the paired end LQ-ssCAGE, Read1 was the Cap-trapped 5’ reads from the transcript start site and Read2 was the sample-barcode sequence and random priming site read. Using only read1, we retained a total of ∼53.3 million reads (median of 783,709.5 reads; **Figure 1B**). Raw sequence data were processed using the Moirai workflow[25] (**METHODS**). The cap-trapped 5′ tags are mapped to the *G. mellonella* genome assembly ASM364042v2 and a high mapping rate (76% of total reads are mapped to reference genome) was obtained after trimming the sequence adapters and primer barcode reads (4%) (**Figure 1C**).

**Figure 1.**
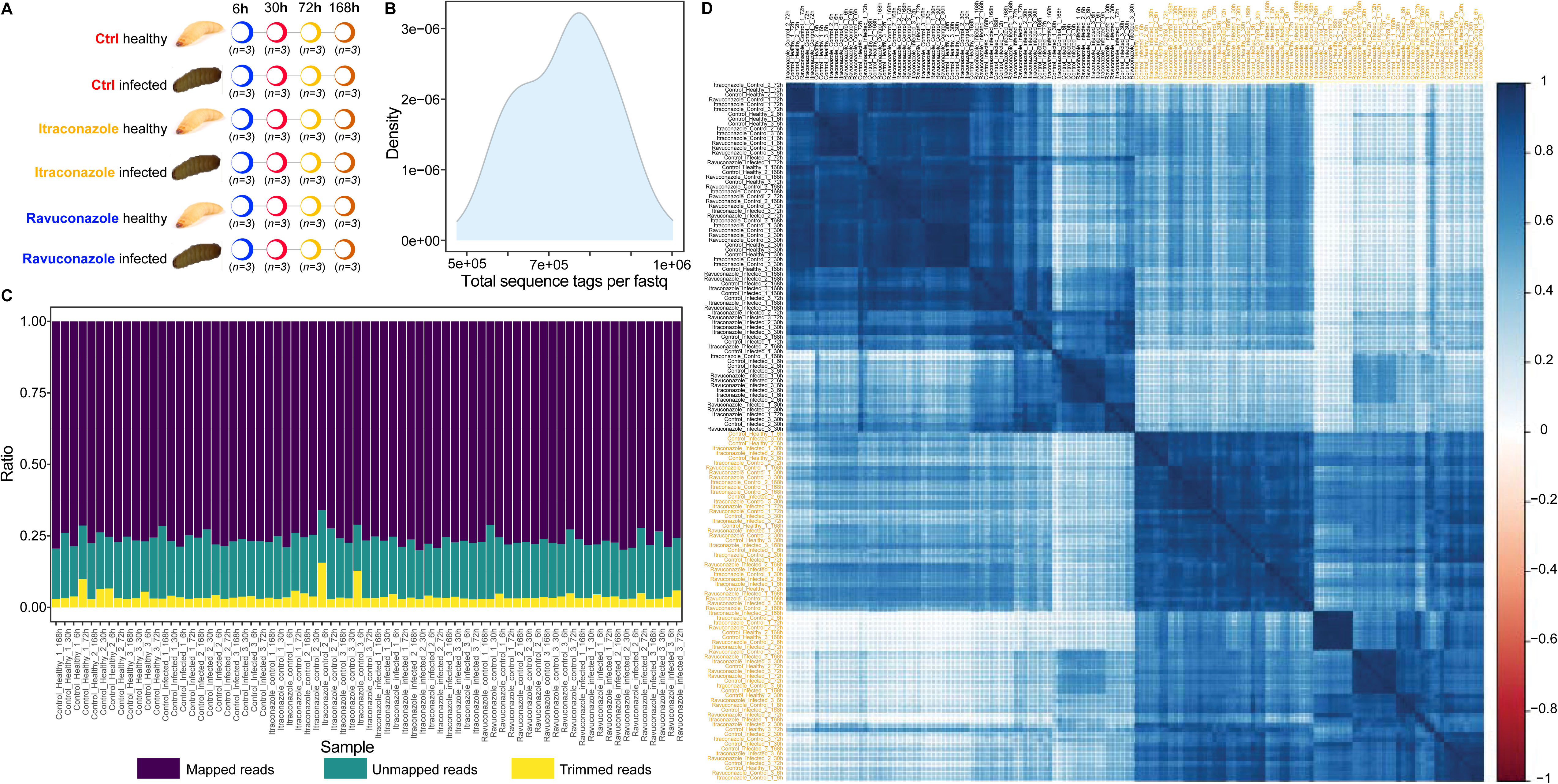
*Galleria mellonella* larvae samples preparation and the quality of the LQ-ssCAGE. **A.** The *Galleria mellonella* larvae were infected with the pathogen *Madurella mycetomatis* (strain mm55) and treated with either Itraconazole or Ravuconazole antifungal. Three groups of samples are generated. The first is control healthy (Ctrl healthy) and control infected (Ctrl infected). The second group, infected and healthy larvae treated with itraconazole (Itraconazole healthy or Itraconazole infected), and the third group, larvae treated with ravuconazole (Ravuconazole healthy or Ravuconazole infected). From each sample group, we generated three replicates and extracted total RNA for LQ-ssCAGE (n = 72) and RNA-seq (n = 72). **B.** Density distribution of the LQ-ssCAGE total sequence tag calculated from the Fastq files. In total, we obtained 53,356,962 sequence tags aligned to the *G. mellonella* reference genome assembly ASM364042v2. **C.** Quality of the mapping of total reads: 4.1% of the reads are trimmed, 19.8% of reads are unmapped, and 76.2% of reads are mapped to the reference genome. **D.** Pair-wise correlation heatmap of the total read count of RNA-seq and LQ-ssCAGE libraries. LQ-ssCAGE samples (orange labels) and RNA-seq samples (black labels). A positive correlation was detected between all samples.

RNA-seq libraries were used to confirm the quality of the LQ-ssCAGE reads and coverage. We performed a pairwise correlation analysis between the total read counts of the mapped RNA-seq reads and the total read counts of the mapped cap-trapped 5′ end tags. **Figure 1D** shows a positive pairwise Pearson correlation between RNA-seq and corresponding LQ-ssCAGE samples.

### TSS and enhancer atlas for G. mellonella

Mapped CAGE tag 5’ ends in BAM format were converted to CTSSs (**METHODS**). CTSSs are the number of CAGE-tag 5’ end mapped to genomic positions on the reference genome. To identify clusters of CTSSs that correspond to the activity of individual TSSs and enhancers, CTSS files are used as input. The R Bioconductor package CAGEfightR[26] was used for the analysis of LQ-ssCAGE and the detailed workflow for the analysis of the LQ-ssCAGE dataset with CAGEfightR is illustrated in (**Figure 2A**). The function quantifyCTSS summarized the number of CAGE tags for each CTSS per sample. The analysis of CTSS usage across all samples (*n*=72) results in a total of 1,208,696 CAGE tags. We subset the raw CTSSs by discarding CTSSs expressed in a single sample (we keep biologically relevant CTSSs), so the final set of CTSSs after filtering was 460,430. The quantified CTSS is stored as a RangedSummarizedExperiment object[27]. This is a 460,430 X 72 matrix with each row in this matrix corresponding to a CTSS (single bp), the object has two rawData names score, which represents the pooled expression, and support which represents the number of samples expressing a feature above zero. The total score and total support were 70,784,739 and 5,188,408 respectively. CTSS is provided as an R rds object (**Data Access**). After identifying the final set of CTSSs, we used this to identify the TSSs and enhancers. In CAGE data, the TSSs can be identified from the uni-directional clusters, while the enhancers are identified from the bi-directional clusters. To find the uni – and bi-directional clusters, pooled CTSSs raw counts were normalized in each sample using the commonly used CAGE data normalization technique Tags-Per-Million (TPM). Secondly, all TPM values were summed across samples. The updated CTSSs object with pooled TPM values were used to locate and quantify the unidirectional clusters known as tag clusters (TCs). The CAGEfightR function quickTSSs() takes CTSSs as input and produces 90,528 TSS candidates. Likewise, the function quickEnhancers() generated 961 bidirectional clusters (candidate enhancers). To obtain biologically relevant TSSs, the raw TCs were further filtered, and only TSSs expressed in at least four samples were kept, this resulted in the final set of 39,410 TSS peaks. The bi-directional clusters were filtered as well. We kept only the exonic clusters (intergenic and intron), which resulted in the final set of 249 enhancers. Finally, we used the combineClusters() function to combine TSSs and enhancers. If TSS and enhancer clusters overlapped, the enhancers alone were kept. The final merged object of TSSs and enhancers contained a total of 39,233. TSS peaks and enhances are provided in different formats (**Data Access**).

**Figure 2.**
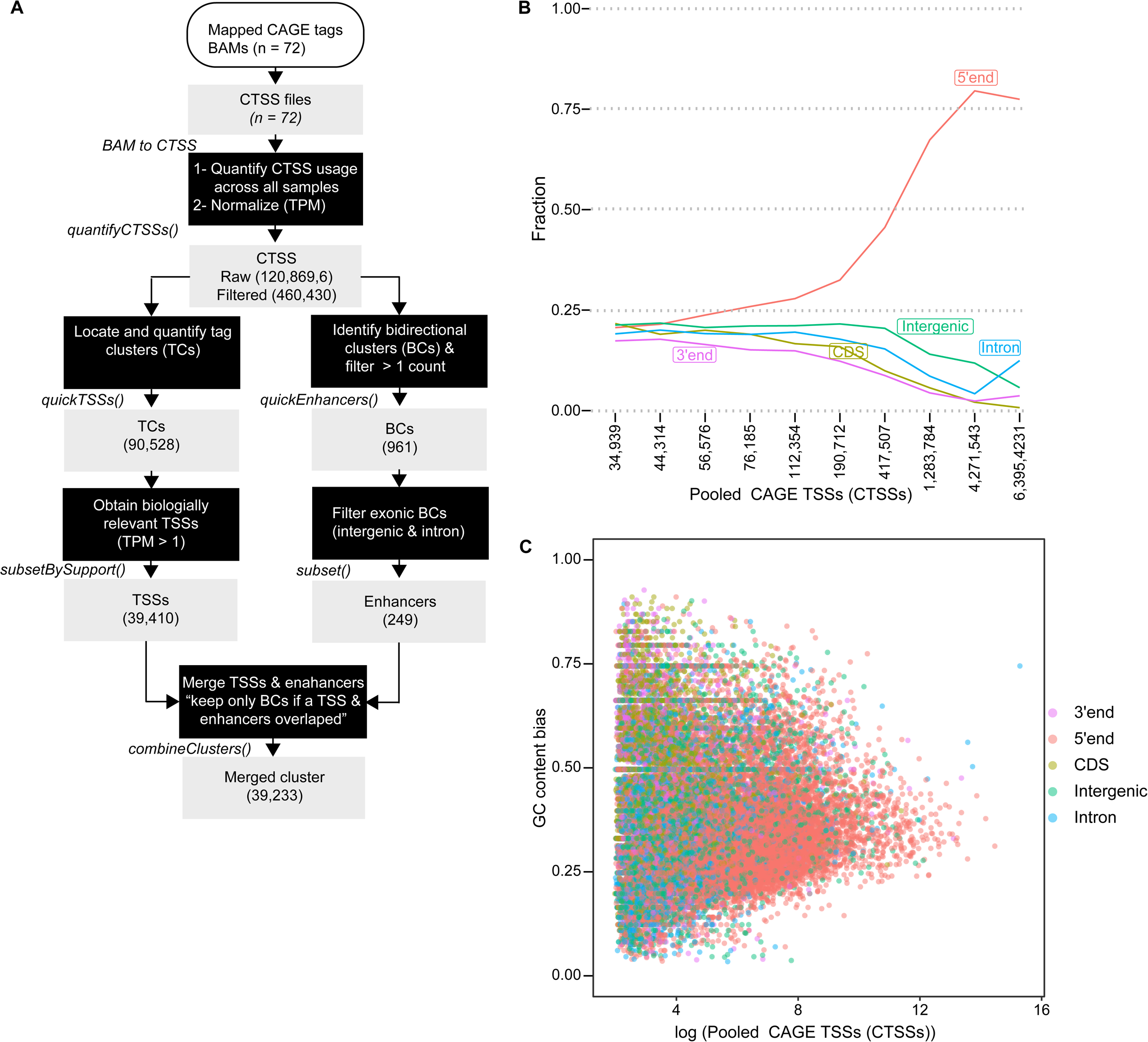
Basic workflow of the LQ-ssCAGE data and characteristics of CAGE TSSs. **A.** Mapped CAGE tags in BAM format (n = 72) are converted to CTSS files. The CTSSs are used to quantify CTSS usage across all samples. A total of 460,430 CTSSs are returned after filtering. The CTSSs are used to locate and quantify the tag clusters (TCs - the candidate TSSs) and identify bidirectional clusters (BCs – candidate enhancers). A total of 90,528 and 961 TCs and BCs, respectively, are identified. To obtain biologically relevant TSSs, only TCs with TPM > 1 are kept, which leads to a total of 39,410 TSSs. To obtain the final list of enhancers, BCs are filtered, and only exonic (intergenic and intron) BCs are kept. The list of TSSs and Enhancers are merged to get a merged cluster totaling 39,233. R functions used at each step are listed. **B.** Pooled CAGE TSSs (CTSSs) intersection with five genomic features. The Y-axis represents the fractions (0 ∼ 1) of CAGE peaks overlapping each of the genomic features (5′end, 3′end, CDS, intergenic and intron). The X-axis shows the number of CTSSs (ordered from highest to lowest). The 5′end genome feature overlaps the top-ranked CTSSs compared to intergenic and 3′end (bottom of the rank list of CTSSs). **C.** GC bias. The scatter plot shows the GC bias (as computed by chromVAR R Bioconductor package) among the genomic features. The 5′end regions have a GC bias of about 30%. The other features have either high levels of GC bias or very low levels of GC bias.

### Cluster peaks with the highest pooled CTSSs overlapping the 5’ end UTRs

The merged object of the TSS and enhancer peaks were annotated by the transcript model of the genome assembly of *G. mellonella*. We created a TxDB object using GenomicFeature package from R Bioconductor. The TxDB consists of 21,263 transcripts. We assigned transcript types to each of the clusters in the merged object, then, we looked at the fraction of the pooled CTSSs that overlap each of the five genomic annotations (**Figure 2B**). We found that the highest pooled CTSS peaks overlap with the 5’ end of the transcript, while only a small number of peaks overlap with the 3’ end, CDS, intergenic, and intronic regions.

### Confirmation of the (di)nucleotide composition of the TSSs

Previous studies in eukaryote and prokaryote core promoters show that core promoters overlap with CpG islands[28] . To validate and confirm the (di)nucleotide composition of the TSSs, we used the addGCBias() function from the chromVAR R Bioconductor package. We found that the mean GC content for the TSS peaks overlapping the 5’ end regions is equal (0.35:1.00) (**Figure 2C**). This analysis shows that LQ-ssCAGE-identified TSS peaks align with known qualities of prokaryote core promoters. Unfortunately, we couldn’t find close moth to the *G. mellonella* (e.g. *Bombyx mori*) with such annotation to compare with our data.

### Identified TSSs and active enhancers confirm known annotation and expression patterns

Annotation with genomic features aims to assign features and functions to unannotated genomic regions. We annotated the merged clusters of TSSs and enhancers (39,233) using the TxDB genomic features class (**METHODS**). We found that many of the TSS clusters (∼29.4 %) overlap the promoter regions of the transcript , whereas the enhancer clusters overlap intronic and intergenic regions only (**Figure 3A**). Looking into the cluster expression, TSS clusters annotated as promoter regions are highly expressed. Novel TSS clusters, not annotated as a promoter, have low expression values except for the ones that are annotated as 5′- UTR and exons (**Figure 3B**). Enhancer clusters annotated as intron and intergenic regions are lowly expressed compared to TSS clusters (**Figure 3B**).

**Figure 3.**
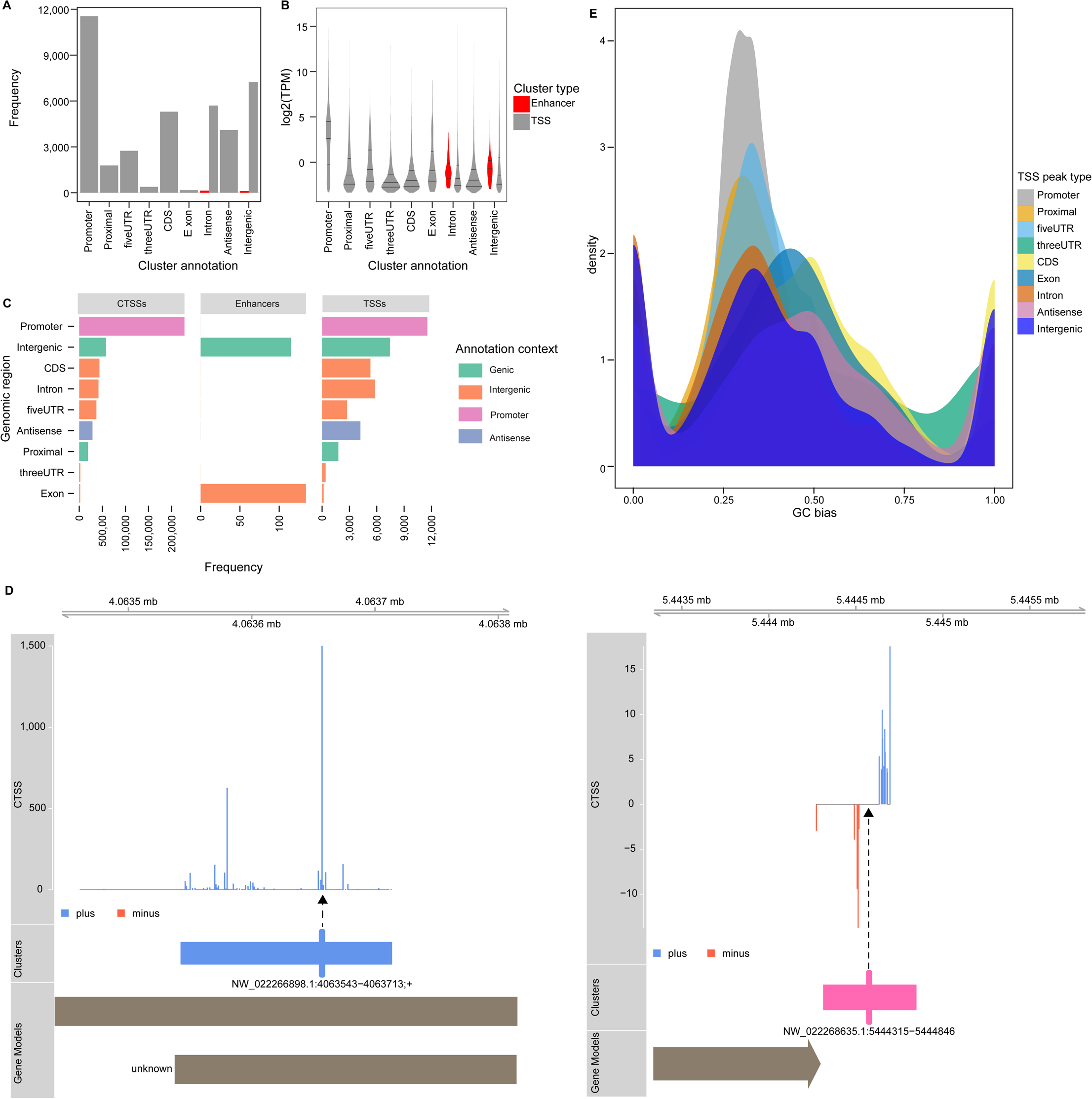
Genomic annotation and expression of the TSS and enhancers peaks and the GC bias of TSS peak type. **A.** Annotation of the cluster by genomic feature categories. Many of the TSS clusters are promoters (11,545/39,233). **B.** Gene expression per cluster annotation. As expected, clusters overlapping promoters have higher gene expression values (log2 TPM) compared to enhancers. **C.** Annotation by the genomic context of CTSSs, enhancers, and TSSs. The majority of CTSSs and TSSs peaks overlap promoter regions, while enhancer peaks overlap genic and intergenic regions. **D.** Genome browser view of TSS peaks and enhancer. The genome browser view consists of three tracks. The top track shows the pooled CTSS signal and the middle track (clusters) shows the identified TC. The gene model track at the bottom of the view shows the known/unknown transcripts model in the genome. Genome browser view of TSS (left panel). The vertical thick bar in the cluster indicates the most used CTSSs within the NW_022266898.1:4063543−4063713;+ cluster. The dashed arrows connect the tick bar to the TSS peak with the most used CTSS. Genome browser view of bidirectional cluster (enhancer NW_022268635.1:5444315−5444846) (right panel). The vertical thick line in the cluster indicates the maximally balanced point within the cluster. The view shows the main characteristic of the active enhancer (the bidirectional pattern of the expression in both the plus and minus strands). **E.** Density distribution of the GC bias of the TSS peaks as obtained by chromVAR function. The peak shows that promoter TSS peak type has a high GC bias.

To confirm the genomic annotation of the CTSSs, enhancers, and TSSs we performed additional annotation by performing annotation by genomic context i.e. genic, intergenic, promoter or antisense. The top annotation category of our CTSSs was promoter, with 49.5 %, 12.6 % of CTSSs were intergenic regions (**Figure 3C**). The rest of the CTSS regions were annotated as CDS, intron, 5′-UTR, antisense, proximal, 3′-UTR or exon. Like the CTSSs, 29.3 % of TSSs were annotated as promoters, and 18.9% were annotated as intergenic (**Figure 3C**). Enhancer clusters were annotated as either genic or intergenic regions only (**Figure 3C**).

**Figure 3D** shows examples of TSSs, and active enhancer clusters as illustrated in the Genome browser view. TSS (left panel) has a high number of pooled CTSS signals compared to active enhancers (right panel). The cluster track of TSS shows the uni-directional (‘+’ strand) expression pattern characteristics of the identified TSS, while the cluster track of the enhancer shows the bi-directional expression characteristic of active enhancers (both strands).

We tested the GC bias of each of the TSS peak types, we found that promoter TSS peaks had elevated GC content (∼0.34: 1.0). The GC bias confirmed the known (di)nucleotide composition of the TSS (**Figure 3 E**).

### Promoters shape and alternative usage of the TSS

Promoters identified by CAGE analysis have been classified into two categories[29], broad or sharp promoters. To classify TSSs, we first calculated the interquartile range (IQR). The distribution of IQRs was bimodal (**Figure 4A**) and showed that most of the TSSs were above or below 10 bp (IQR). We used 10 bp IQR as the cut-off value to classify the TSSs into broad or sharp promoters, the classification resulted in 4,489 broad promoters and 1,818 sharp promoters. **Figure 4B** demonstrates two examples of broad (left) and sharp (right) promoters. The broad promoter “NW_022266059.1:3349304−3349414; +” had multiple spaced TSSs, while the sharp promoter “NW_022266059.1:4929965−4930023; +” had a single well defined TSS as described by Haberle and Stark[30]. We determined the expression difference between broad and sharp promoters and noted slight differences in the expression between the two categories (**Figure 4C**). Next, we looked at differential TSS usage. Differential TSS usage aims to analyze if a gene uses different TSSs under different conditions, analogous to alternative splicing identification in RNA-sequencing[31]. The edgeR[32] function diffSpliceDGE() tests for differential exon usage between experimental conditions. We used diffSpliceDGE() to test for differential TSS usage instead of exon. It tests if a TSS reveals similar changes to other TSSs in a particular gene or not. Our analysis of the TSS usage by diffSpliceDGE() indicated that the majority of the genes used a single TSS, while only a few genes used more than one TSS to initiate the expression of the gene (**Figure 4D**).

**Figure 4.**
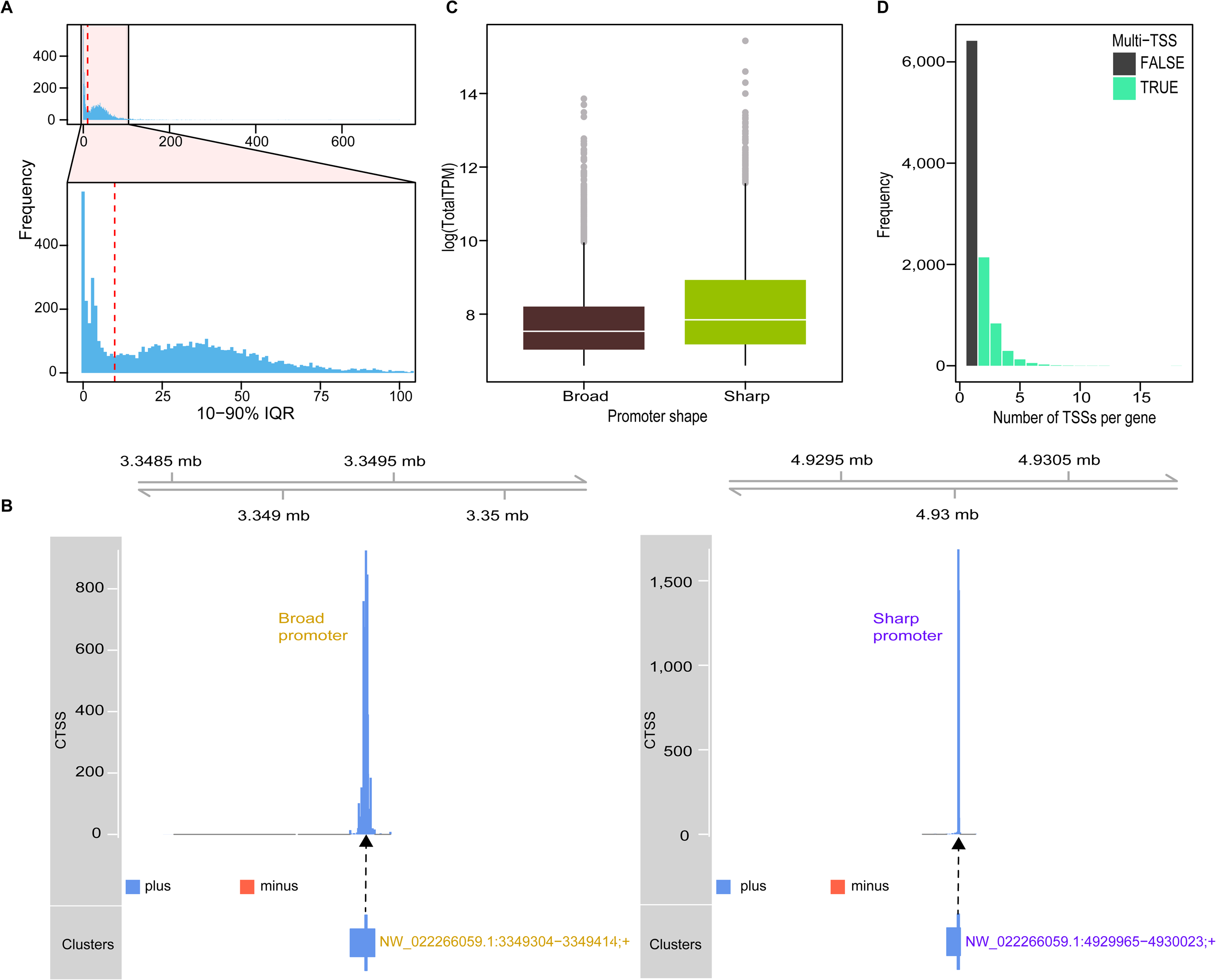
TSSs shape analysis and TSS gene usage **A.** The bar plot shows the distribution (bimodal) of the Interquartile Range (IQR). IQR divides the TSS into sharp and broad TSSs. The dashed line indicates that TSSs are either below or above an IQR of ten. Broad promotors have IQRs below ten, and sharp promoters have IQRs above 90. **B.** Promoter shapes. An example of a broad promoter (left panel) with multiple spaced TSSs, and a sharp promoter (right panel) with a single TSS. **C**. Distribution of expression between broad and sharp promoters shows a slight difference in the normalized expression between the two classes of promoters. **D.** TSS usage by gene. Differential TSS usage tests whether a gene uses alternative TSSs under different conditions. The figure indicates that the majority (6,415/9,903) of *G. Mellonella* genes utilize a single TSS, while 3,488/9,903 utilize multiple TSSs.

### TSS-enhancers interactions

Genome-wide Chromatin Interaction Analysis with Paired End-Tag sequencing (ChIA-PET) was used to identify promoter-enhancer interactions [33, 34]. Furthermore, CAGE data can be used to predict physical interaction between enhancers and TSSs[23], therefore identifying which genes are regulated by enhancers.

In CAGE data, TSS and enhancers being in proximity and having high correlation expression, are candidates for TSS-enhancer links or interactions. Using findLinks() function we calculated pairwise correlations in expression (TPM) between TSSs and enhancers within a distance of 50 bp with Kendall’s tau. Considering positive correlation only and *p-value* < 0.05, we found that 633/39,410 TSSs are interacting with 181/249 active enhancers. **Figure 5A** shows that the enhancer “NW_022270367.1:4181104−4181512” has multiple interactions as shown by the links track. The TSS “NW_022270367.1:4182931−4183061; +” has the highest correlation upstream of the gene LOC113521978 (*p* = 0.00022). The list of TSS-enhancer links is provided as rds R object (**Data Access**). We compared the average CAGE expression between interacting and non-interacting TSSs and observed that the interacting TSSs have higher average gene expression. The mean of the expression (TPM) between the two classes of TSSs was 10,202 and 1,684 for interacting and not interacting TSS, respectively. (**Figure 5B**). The highly correlated interactions are provided as an R rds object (**Data Access**).

**Figure 5.**
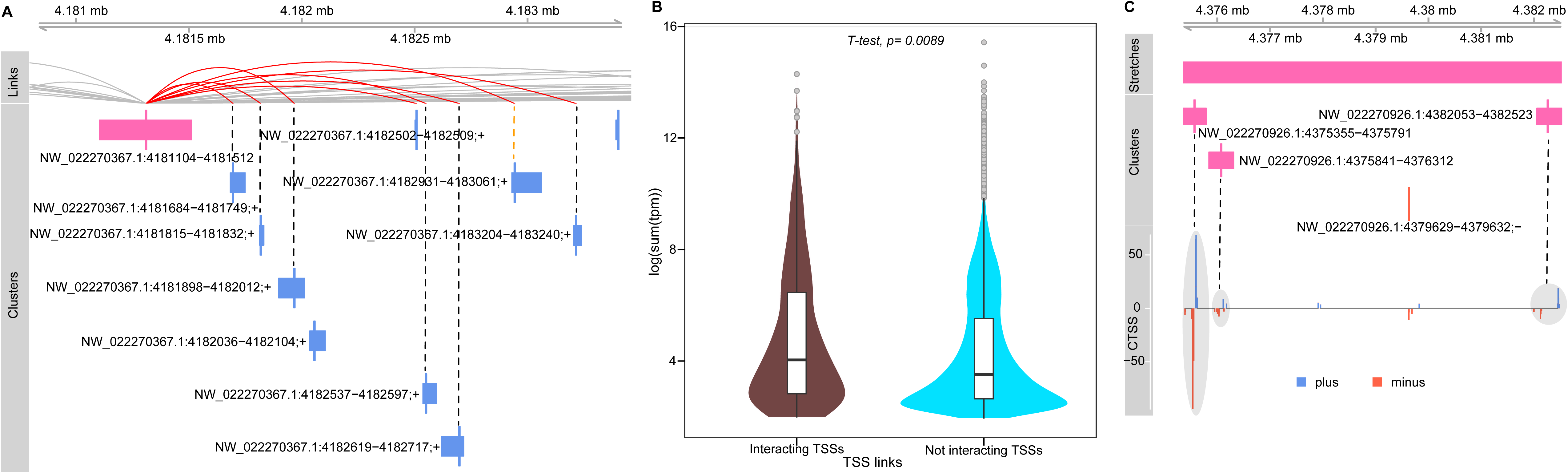
TSS-enhancer interactions and stretched enhancers. **A.** Genome browser view of interacting TSSs and enhancers. The genome view consists of three tracks: links, identified clusters, and pooled CTSS signals. By using the distance and expression correlation patterns between TSSs and enhancers, we predict enhancers-TSSs links. The links track shows the strength of correlation between 8 TSSs and the NW_022270367.1:4181104−4181512 enhancer. The highest correlation was found between NW_022270367.1:4182931−4183061;+ TSS and the enhancer (dashed orange line). **B.** Variations in the gene expression between interacting and non-interacting TSSs. The violin plot shows that TSSs interacting with enhancers are highly expressed compared with the non-interacting TSSs. **C.** Genome browser view of super-enhancers. Super or stretch enhancers are closely spaced enhancers. We found two stretched enhancers. The genome browser view shows stretch consists of three enhancers with bidirectional expressions. The stretches are visualized by the Stretches track on top of the genome browser view.

### Super enhancers

Super-enhancers (stretch enhancers) are “*groups of putative enhancers in close genomic proximity with unusually high levels of mediator binding, as measured by (ChIP-seq)*” [35] . Compared to singleton enhancers, stretch enhancers show different functions and chromatin characteristics[35].

The function findStretches() allows the identification of stretches by grouping nearby proximal enhancers into groups, each consisting of a cluster of enhancers. After identifying the nearby groups, an average pairwise correlation between them was calculated.

In our data, we identified two stretches each having 3 bi-directional clusters and an average correlation of 0.06 and 0.09. **Figure 5C** shows the first stretch with three bi-directional enhancers overlapping intergenic regions. The identified stretched are provided as R rds object (**Data Access**).

*Homer de-novo Motif for G.mellonella and TF expression analysis*

HOMER[36] command findMotifsGenome.pl was used to predict the enriched motifs and corresponding TFs for *G. mellonella* using TSS peak regions from LQ-ssCAGE. Two types of TFs are predicted (**Supplementary Information**) the Homer de novo motif and Homer known motif enrichment results. The de-novo motif prediction results in 27 motifs with a *p-value* < 0.0001 (false positives were excluded). We selected the top-ranked motifs for expression analysis (**Figure 6**). The motifs are ordered by log *P-value*. For each motif, we obtained the TF from the Jaspar database[37], since *G. mellonella* was not supported by Jaspar, we obtained the ortholog for each TF using NCBI blastx[38]. LQ-ssCAGE gene expression of the predicted TFs shows a difference in gene expression between healthy and infected samples (broadly expressed). The top-ranked TF was Zinc finger protein PLAG1-like (LOC113511025), it has a biological process GO term of regulation of gene expression.

**Figure 6.**
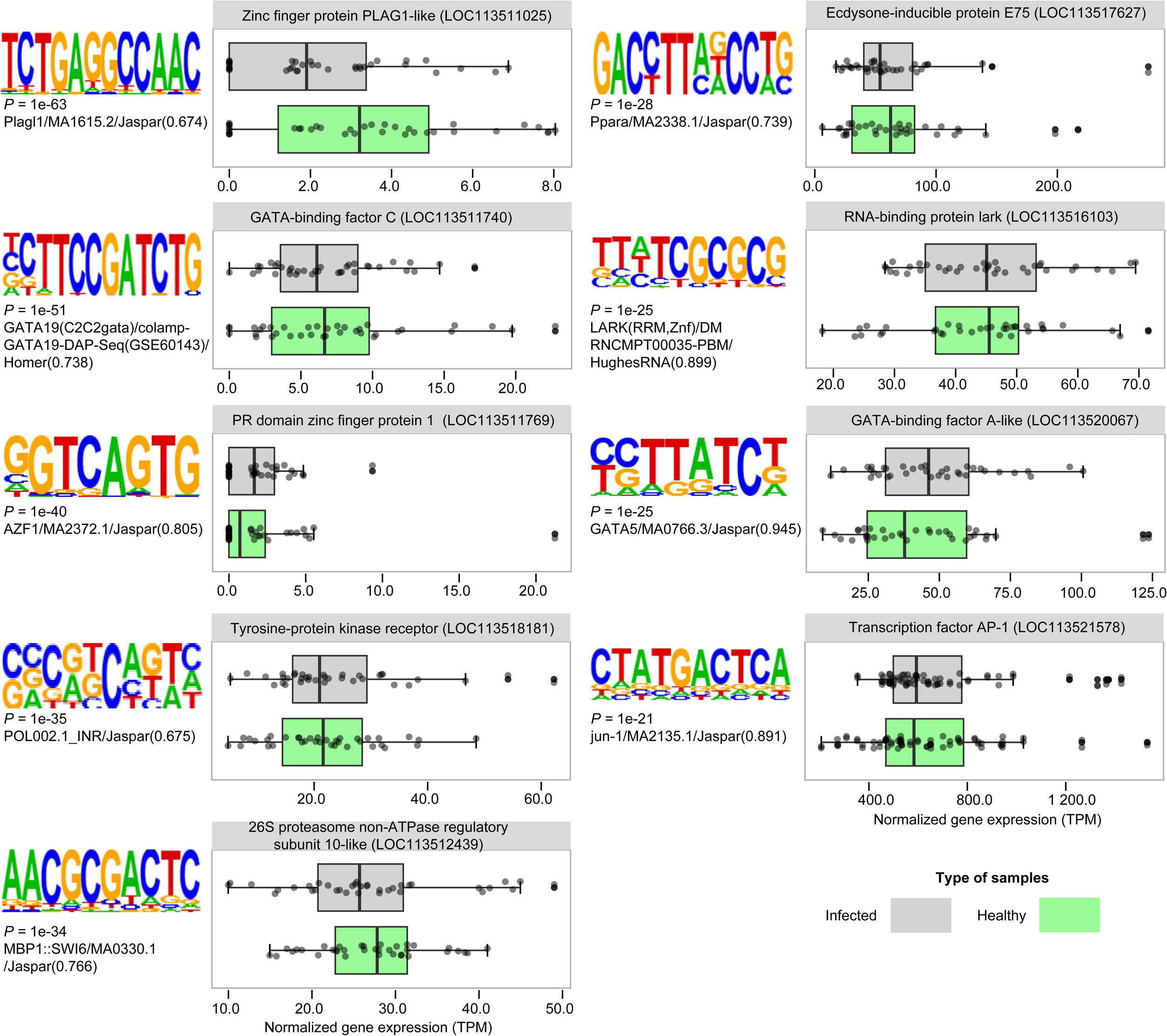
Motifs prediction and analysis; The list of TSS peaks is used as input for HOMER suite for de novo motif prediction. The figure shows the top predicted motifs and their corresponding *G. mellonella* transcription factor expression, each box plot shows the CAGE expression as TPM for the newly identified top TFs across samples between infected and healthy *G. Mellonella*.

### Genomic views of the G. mellonella annotation on ZENBU genome browser

Genome browsers are a useful tool to visualize and explore genomic features in a 2D view using genomic coordinates of the genes (features). To understand the localization of the mapped TSS peaks and identified enhancers. We first uploaded the genome assembly (ASM364042v2) to the ZENBU genome browser. All annotated features in ASM364042v2 are searchable and can be explored (e.g. zoom-in or out). On top of the genome assembly, the LQ-ssCAGE mapped reads are also uploaded to the genome browser. Together with mapped CAGE tags (in BAM format), detailed metadata about each sample is uploaded as well. The metadata, together with mapped CAGE tags, allows grouping of the sample experiments per treatment (timepoint, replication, etc.) and makes for a more interesting ZENBU genome browser view.

Based on our experimental design, we created four tracks with different groupings based on the experiment’s conditions. https://fantom.gsc.riken.jp/zenbu/gLyphs/#config=Galleria_mellonella_TSS.

**Figure 7** shows the CAGE expression of all samples. Infected samples have higher CAGE expression compared to healthy samples. Both in the +/- strand for the genomic region “NW_022266059.1:12016449..12066485+” (**Figure 7**).

**Figure 7.**
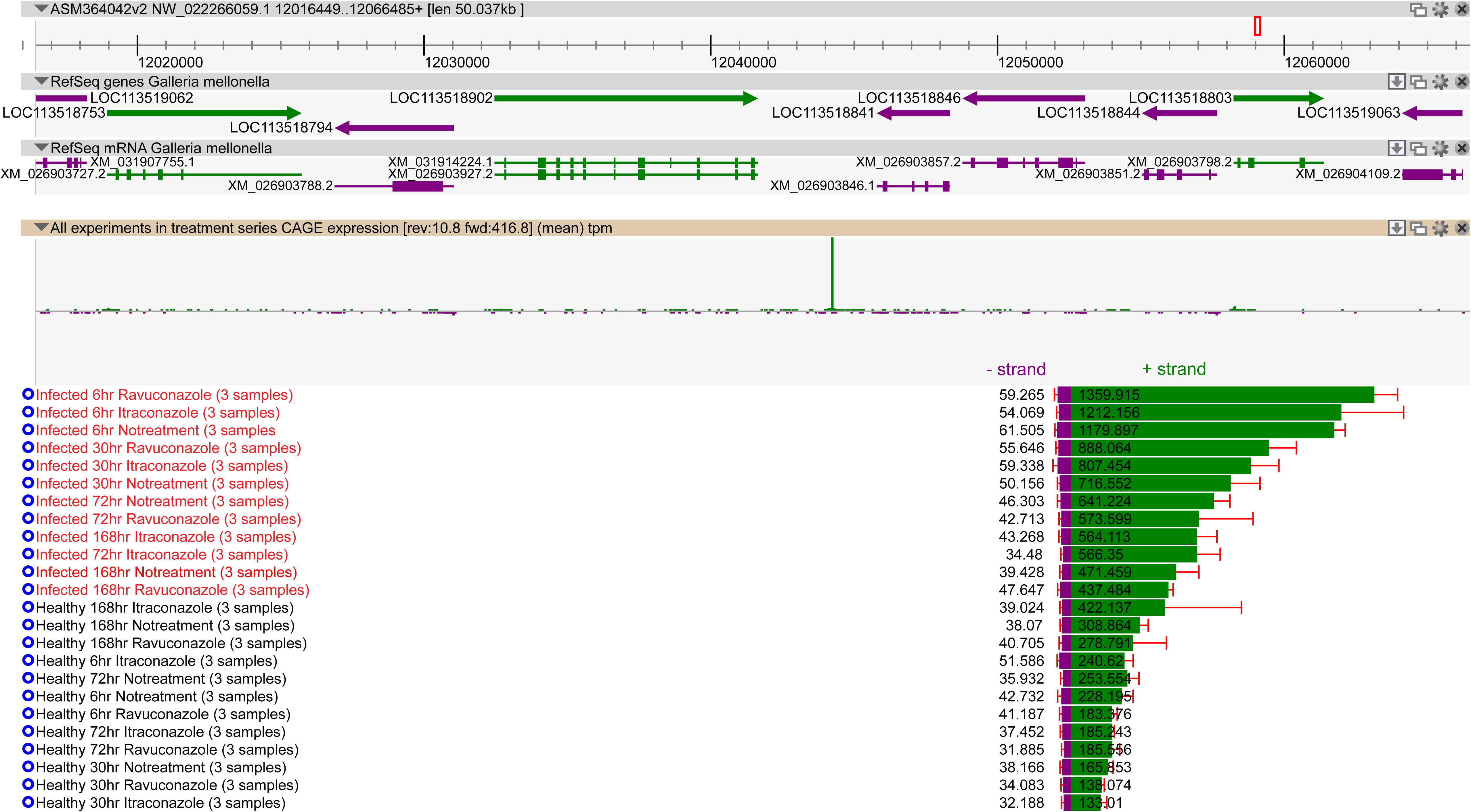
ZENBU genome browser view of TSS region for all samples. The top panel shows the refSeq genes for *G. mellonella* and the bottom panel shows CAGE expression in both strands. Samples are grouped by experimental conditions, either infected or healthy.

## DISCUSSION

The lack of TSS information on the invertebrate *G. mellonella* genome will limit its utilization as an experimental model in the wider research community. To address this, we performed genome-wide mapping and analysis of TSSs in *G. mellonella* larvae using LQ-ssCAGE. LQ-ssCAGE libraries (n=72) were sampled under differential conditions and provided us with a comprehensive set of CTSSs. The comparison of the mapped 5’ end tags and RNA-seq reads shows a high correlation between the two methods and confirms the reproducibility of the expression profiles by two independent sequencing methods, as reported by Kawaji *et al* [39].

LQ-ssCAGE allowed us to map 460,430 CTSSs for *G.mellonella* larvae. The quality of the resulting CTSS library was confirmed by genomic annotation and GC bias. The annotation of the CTSSs confirmed a promoter-rich CAGE profile and the analysis of GC bias confirmed a highly elevated GC level. TSS regions annotated as promoters are highly expressed compared to enhancers, the variation of the expression patterns between promoters and enhancers here lines up with previously reported results[40].

From the CTSS profile, we identified 39,410 TSS and 249 active enhancers by their characteristic uni-directional or bi-directional peaks respectively, we used these characteristics to further assign genomic annotations as TSS and enhancer.

Genome views of the TSS peaks and enhancers confirm the expression directionality of each type of cluster and showed that TSS peaks are annotated as promoters while the enhancers are annotated as intergenic and genic.

The promoter shape plays a fundamental role in the gene expression regulation in animals[29, 30, 41]. To characterize the TSS shape, we computed the IQR. Before computing IQR, we filtered lowly expressed TSSs and instead, we used only highly expressed TSSs with an average TPM > 10. TSSs with low expression have low width and low number of CAGE tags therefore, they can’t show the variations in the shape[19]. Taking 10% pb IQR as the cutoff, we divided the TSS into broad (n=4,489) and sharp (n=1,818) promoters. The two classes of TSS show different architectures in the genome browser view and show variations in CAGE gene expression. In addition to the promoter shape, we analyzed the differentiation of TSS usage (i.e. genes use different numbers of TSSs under different conditions)[19]. We show that in *G. mellonella*, most of the genes use a single TSS, but some genes use more than one TSS to initiate expression.

One of the features of CAGE is that it enables the prediction of the physical interaction between enhancers and TSSs. This kind of interaction indicates which gene is regulated by which enhancer. As reported previously, TSSs and enhancers in proximity and have highly correlated gene expression and are potentially interacting[23]. We identified a set of interactions between TSS and enhancers. This enabled us to classify the TSS as either interacting TSSs or not interacting TSSs. We observed quite different gene expressions between the two classes of the TSSs.

As reported by Pot and Lieb[35], super-enhancers (stretched enhancers) demonstrate chromatin characteristics and function differently from singleton enhancers. In our LQ-ssCAGE data, we detected two stretches, each with 3 clusters of enhancers (9 super enhancers).

To compare our TSS annotation of *G. mellonella*, we investigated the availability of TSS annotations for closely related species. First, we considered *Bombyx mori* [NCBI taxonomy: 7091] from the family Lepidoptera and same clade (Obtectomera) as *G. mellonella*. In SilkBase[42] the only TSS set was obtained computationally by mapping the BmN4-derived TSS-tags to the *B. mori* genome[43]. With no characterization of TSS and promoters.

## Conclusions

Our results contribute to the genome annotation of *G. mellonella* and will enable the study of gene regulation research in this unique model. The raw sequence reads, and processed CAGE data are available via NCBI GEO[44] and ZENBU genome browser [45]. The resulting TSS atlas in this study, will be a centerpiece for the development of a reference transcription start site for the *G. mellonella like the* human and mouse refTSS we developed previously (https://reftss.riken.jp/)[9].

## METHODS

*G. mellonella* Acquisition and infection

*G. mellonella* larvae were obtained from Forelshop (Tremelo, Belgium) and kept in the dark on wood shavings at room temperature until use. Within five days of receipt, larvae of approximately 300-500 mg were selected for experimental use. The selected larvae were divided into Petri dishes containing 90 mm Whatman filter paper and five larvae per dish.

An inoculum of *M. mycetomatis* strain MM55 was prepared following the method described by Kloezen et al[2] . In short, the mycelium was harvested, transferred to RPMI-media supplemented with 0.35 g/L L-glutamine, 1.98 mM 4- morpholinepropanesulfonic acid (MOPS) and 100 mg/L chloramphenicol, sonicated for 30 seconds at 10 microns (Soniprep 150 plus, MSE, The United Kingdom) and incubated for two weeks at 37 C. Next, the mycelium was harvested using vacuum filtration (Nalgene, Abcoude, the Netherlands, using a 22-micron filter (Whatman)). The fungal biomass was scraped from the filter, and the wet weight was measured before resuspending the mycelium in phosphate-buffered saline (PBS). The obtained suspension was sonicated for 120 seconds at 10 microns, washed in PBS, and diluted to a final concentration of 4 mg per 40 µL. The larvae were infected using the prepared inoculum by injection in the last left proleg using an insulin 29G U-100 needle (BD Diagnostics, Sparsk, USA). After 4, 30, and 52 hours post-infection, larvae were treated with itraconazole or ravuconazole to a final concentration of 5.71 mg/kg, a clinically relevant dose of 400 mg based on an average human with a weight of 70 kg[46]. The course of the infection was monitored by infecting and treating separate groups consisting of 15 larvae per group, and survival was recorded daily for ten days. During all experiments, Pupae were removed from the equation, and non-infected larvae were included as a control.

### RNA isolation

RNA was isolated at 6-, 30-, 72- and 168 hours post-infection. The contents of three larvae were pooled, flash-frozen with liquid nitrogen, and mechanically crushed using a pestle and mortar. The resulting powder was suspended in RLT buffer (supplemented with 1% β-Mercaptoethanol), provided in the RNeasy Mini Kit (Qiagen, Germany). The samples were incubated at 57 C for 3 minutes before proceeding according to the manufacturer’s instructions.

### RNA-Seq library preparation and sequencing

RNA-seq library was prepared using MGIEasy RNA Directional Library Prep Set (MGI Tech) according to the manufacturer’s protocols. We used 1 μg of total RNA for each sample for library preparation. PolyA+ RNA was enriched from the total RNA with oligo dT beads before preparing the RNA-Seq library. Resulting RNA library was QCed using Qubit dsDNA HS Assay Kits (Thermo Fisher Scientific) and Agilent TapeStation, with High Sensitivity D5000 Reagents (Agilent Technologies), and sequenced with 100 bp paired end reads using a DNBSEQ G400 Sequencing platform (MGI Tech).

### LQ-ssCAGE library preparation and sequencing

LQ-ssCAGE started with Reverse Transcription (RT) of 200 ng of total RNA to generate 5’ end of the capped RNA/first-strand cDNA hybrids. The RNA/cDNA hybrids were cap-trapped and adapted with illumina sequencer-compatible barcode-tagged adaptors[18]. No PCR amplification was performed during the LQ-ssCAGE library preparation. The number of resulting libraries was checked by qPCR using KAPA Library Quantification Kits (Kapa Biosystems). LQ-ssCAGE library sequenced using Illumina HiSeq2500 platform in Rapid mode with Paired-end read type and 50 base read length.

### LQ-ssCAGE data processing

LQ-ssCAGE Paired-end sequence data provides cDNA sequence as Read1 and sample barcode sequence as Read2. To demultiplex sample barcode sequence, an in-house script is used with Read1 and Read2 data. Analysis of LQ-ssCAGE tags is performed by Moirai workflow[25] using Read1 cDNA read. The genome mapping software used was STAR (v2.5.3)[47]. BAM format files are worked on using samtools (v1.14)[48]. Bedtools[49] (v2.30.0) was used for counting LQ-ssCAGE tags (**Supplementary Figure 2**).

### TSS peaks calling

A genome-wide atlas of TSSs was created for *G. mellonella* using LQ-ssCAGE. The mapped TSSs enabled the precise identification of promoters and different types of enhancers (active and super-enhancers). To identify the promoters and enhancers, we created Biostrings-based genome data packages (BSgenome data packages) using BSgenomeForge Bioconductor R package. BSgenome data packages were used by CAGEfightR to identify TSSs and enhancers[19].

CAGEfightR analysis workflow performed at three levels:

1. CTSS-level analysis to analyze the number of CAGE tags per CTSS.
2. Cluster-level analysis takes the results of CTSS-level analysis to identify clusters of CTSSs. It identifies two types of clusters, uni-directional (candidate TSSs) and bi-directional (candidate enhancers)
3. Gene-level analysis takes as input, the set of annotated TSSs to look at the related gene expression.

BSgenome was used for annotation of the CAGE-defined TSSs and gene-level expression. The mapped reads for each library were converted to CTSSs in BED format using an in-house script as described earlier. The output of CTSS-level analysis was the CTSS tag clusters. For each CTSS, counts were summarized and normalized by TPM. The normalized CTSS counts were used to calculate the pooled CTSS, or CTSS signal across all samples. To reduce noise, pooled CTSSs were further processed by filtering out single tags spread across the genome. The filtering was accomplished by removing CTSSs detected in only a single sample. The final set of pooled CTSSs were used for the analysis at the level of clusters of CTSSs (Promoters and enhancers identification) and analysis at the level of annotated genes.

### Prediction of the transcription factors binding sites (TFBS)

To predict transcription factor binding sites for *G. Mellonella*, we used findMotifsGenome.pl from Homer[36]. The input for the command is the set of identified promoters and enhancers from LQ-ssCAGE data. Also, the Homer command requires a FAST and GFF file for the *G. mellonella* genome, which was obtained from NCBI genome.

### Creating genomic views of the *G. mellonella* new annotation

To create ZENBU genome browser views, first, we fetched the FASTA format DNA and protein sequence alignment file (.fna) for the genome assembly ASM364042v2. The .fna file was downloaded from: https://ftp.ncbi.nlm.nih.gov/genomes/all/GCF/003/640/425/GCF_003640425.2_ASM364042v2/GCF_003640425.2_ASM364042v2_genomic.fna.gz

Next, we retrieved the annotation files (.gff) for the assembly ASM364042v2 from: https://ftp.ncbi.nlm.nih.gov/genomes/all/GCF/003/640/425/GCF_003640425.2_ASM364042v2/GCF_003640425.2_ASM364042v2_genomic.gff.gz

The uploaded files were checked for completeness by searching randomly for several genes and confirming their annotation. Next, we uploaded the 72 aligned LQ-ssCAGE tag counts (stored in BAM format). Each BAM represents a single sample.

## Declarations

### Ethics approval and consent to participate

NA

### Consent for publication

NA

## Availability of data and materials

Raw and processed dataset associated with this study is available for download from NCBI GEO under the accession number GSE282923. The accession number contains the following files:

Raw sequence reads from LQ-ssCAGE

Read1 cDNA reads fastq files (n=72).

CTSS BED files

The BED file stores the CTSS information. For each fastq file, we deposited the corresponding CTSS file (*.ctss.bed). The BED file contains the genomic coordinates of TSSs and the tag count that supports each TSS. Each row in the BED file represents a 1-based coordinate of the TSS on the genomic locus. The file is organized into six columns as follows:

| Column | Description |
| --- | --- |
| Genomic locus | GeneBank locus / NCBI Reference Sequence e.g. (NW_022279282.1) |
| Start position of the locus | Number |
| End position of the locus | Number |
| Barcode e.g. (LS3892_GATGCAT) | String (not used) |
| Number of CAGE 5' tags | Number of CAGE tag 5' ends mapped to the locus and supporting the TSS |
| Genomic strand e.g. (+ / - _) | + (forward) or - (reverse) strand |

Processed data files from R (R Data Serialization) format

The results from CAGEfightR processing of the CTSS files are deposited in R Data Serialization format (.rds). The function readsRDS() can be used to read and load the .rds object in the R environment. The following files are available:

CTSSs.rds

Contains RangedSummarizedExperimen an output of the quantifyCTSSs() function. The two assays in this object are CTSS raw and normalized (TPM) counts. In addition to the two assays, the object stores the metadata columns (colData). The CTSS object is a 460,430 X 72 matrix.

TSSs.rds

Contains RangedSummarizedExperimen of the identified TSS peaks (uni- directional clusters). TSSs.rds was an output of the following scripts:

TCs <- quickTSSs(CTSSs)

TSSs <- TCs %>% calcTPM() %>% subsetBySupport(inputAssay="TPM", unexpressed=1, minSamples=4)

The two assays in the TSS object are raw and normalized (TPM) counts. In addition to the two assays, the object stores the metadata columns (colData). The TSSs object is a 39,410 X 72 matrix.

Enhancers.rds.gz

Contains RangedSummarizedExperimen of the identified enhancers (bi-directional clusters). Enhancers.rds was an output of the following scripts:

BCs <- quickEnhancers(CTSSs)

BCs <- subsetBySupport(BCs, inputAssay="counts", unexpressed=0, minSamples=4)

Enhancers <- subset(BCs, txType %in% c("intergenic", "intron"))

The two assays in this TSS object are raw and normalized (TPM) counts. In addition to the two assays the object stores the metadata columns (colData). The Enhancers object is a 249 X 72 matrix.

cor_links.rds.gz

This rds stores the GInteractions object with 3574 interactions and 4 metadata columns.

stretches.rds.gz

This rds stores the set of identified super enhancers as GRanges object with 2 ranges and 4 metadata columns.

RawCount.rds.gz

This rds stored the raw expression table for all samples. Each row represents a gene, and each column represents a sample 9903X72.

ZENBU genome browser views

Genome browser views, can be accessed through the link below https://fantom.gsc.riken.jp/zenbu/gLyphs/#config=Galleria_mellonella_TSS

## Supplementary material

The CTSS, TSS peak coordinates, and Enhancers coordinates in Excel format are available for download from the following page.

https://dmsgrdm.riken.jp:5000/9ay8r/

## Authors’ contributions

LQ-ssCAGE experiments, RM, TK. HK., RNA-seq experiments, SN., RNA-seq and LQ-ssCAGE raw data processing MT., AH., IA., sequence data management CT., administration of LQ-ssCAGE and RNA-seq experiments, YO. Write original draft IA., MK., RM., MT., *G. mellonella* sample preparation and RNA extraction, MK., Creation of ZENBU genome browser views, JS., analysis of LQ-ssCAGE IA, results interpretation IA., TK. Visualization, JS., IA., Fund acquisition IA., WdS., TK. Project supervision IA., WdS., TK.

## Acknowledgments

We thank Atsui Hiroto, Nana Tamura, Nobuyuki Takeda, Teruaki Kitakura, and Akira Furukawa for providing technical and administrative support. We thank RIKEN Center for Integrative Medical Sciences sequence platform for their helpful support in sequencing RNA. We thank the Drugs for Neglected Diseases initiative (Geneva) for support of MycetOS.

The authors would like to thank Scott Walker from RIKEN Center for Integrative Medical Sciences for English proofreading.

