## Supplementary figures and images for "Genome-wide mapping of the *Galleria mellonella* larvae transcription start sites during fungal infection and treatment"

### Supplementary_Figure 1

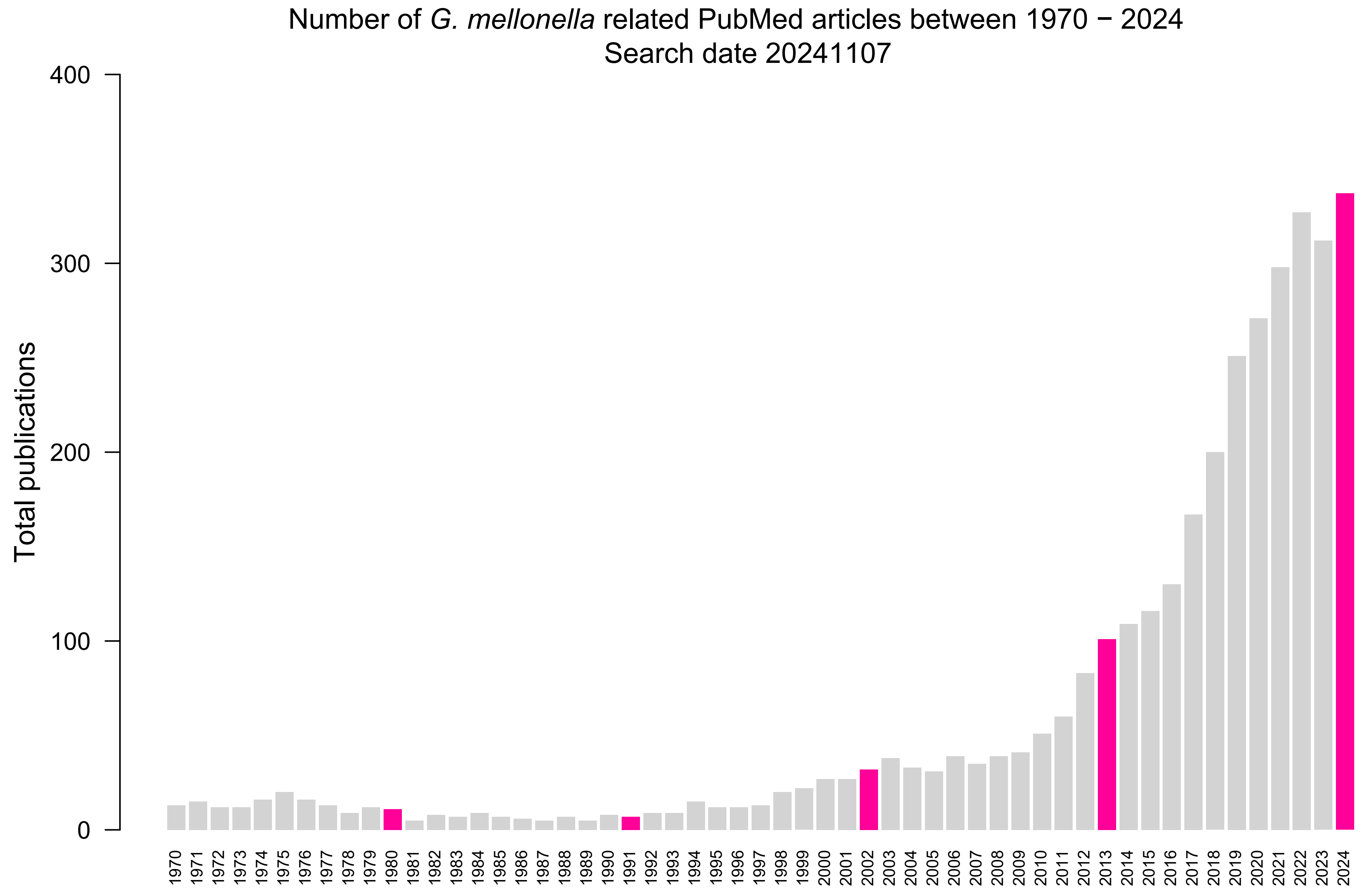

### Supplementary_Figure 2

Supplementary figure 1 LQ-ssCAGE data processing

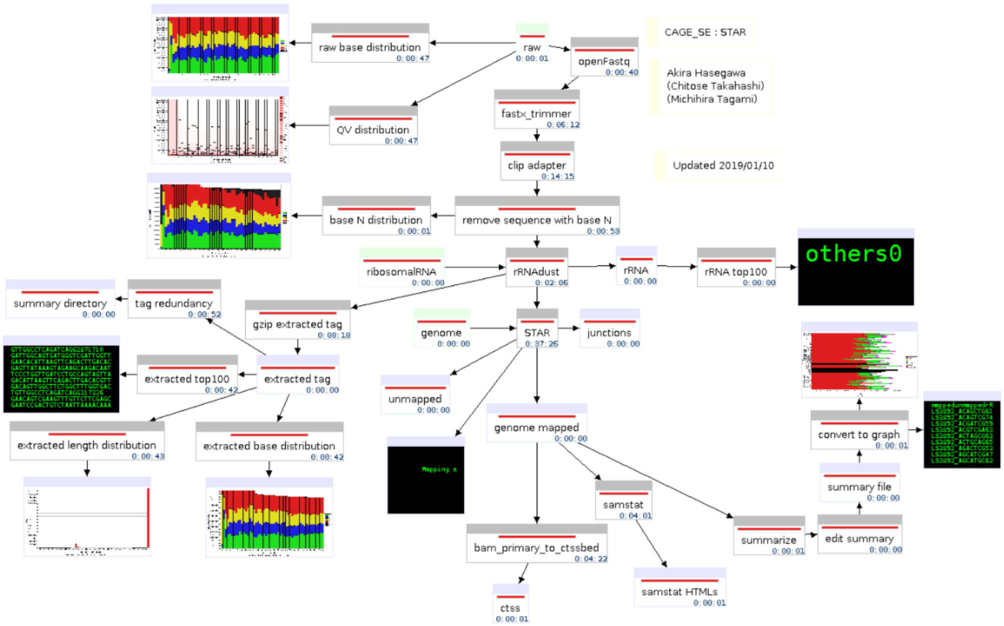
