## Supplementary_Information for "Genome-wide mapping of the *Galleria mellonella* larvae transcription start sites during fungal infection and treatment": homerResults.html

../results/OutputResults// - Homer de novo Motif Results


### Homer *de novo* Motif Results (../results/OutputResults//)

Non-redundant Motif File of Results  
Known Motif Enrichment Results  
Gene Ontology Enrichment Results  
If Homer is having trouble matching a motif to a known motif, try copy/pasting the matrix file into
STAMP  
More information on motif finding results: HOMER
| Description of Results
| Tips
  
Total target sequences = 7192  
Total background sequences = 90822  
\* - possible false positive  

|  |  |  |  |  |  |  |  |  |
| --- | --- | --- | --- | --- | --- | --- | --- | --- |
| Rank | Motif | P-value | log P-pvalue | % of Targets | % of Background | STD(Bg STD) | Best Match/Details | Motif File |
| 1 | A C G T A G T C A C G T C A T G C T G A C A T G A C T G A G T C G T A C C G T A G C T A A G T C | 1e-63 | -1.462e+02 | 0.61% | 0.01% | 6.0bp (49.8bp) | Plagl1/MA1615.2/Jaspar(0.674) More Information \| Similar Motifs Found | motif file (matrix) |
| 2 | A G C T A T G C C G A T A C G T A G T C A G T C A C T G C G T A A C G T A T G C A C G T A T C G | 1e-51 | -1.197e+02 | 0.39% | 0.00% | 22.6bp (39.3bp) | GATA19(C2C2gata)/colamp-GATA19-DAP-Seq(GSE60143)/Homer(0.738) More Information \| Similar Motifs Found | motif file (matrix) |
| 3 | T A C G A C T G A C G T A T G C G T C A T C A G A C G T T C A G | 1e-40 | -9.353e+01 | 39.62% | 32.10% | 51.9bp (64.0bp) | AZF1/MA2372.1/Jaspar(0.805) More Information \| Similar Motifs Found | motif file (matrix) |
| 4 | T C A G T G A C A T G C T C G A C T G A A G C T C T A G G C A T G A T C C T G A | 1e-38 | -8.806e+01 | 21.97% | 16.09% | 55.3bp (63.0bp) | vis/MA0252.2/Jaspar(0.735) More Information \| Similar Motifs Found | motif file (matrix) |
| 5 | A T G C T A G C A T G C C T A G A C G T A G T C G T C A A C T G C G A T G T A C | 1e-35 | -8.223e+01 | 22.39% | 16.64% | 51.1bp (63.1bp) | POL002.1\_INR/Jaspar(0.675) More Information \| Similar Motifs Found | motif file (matrix) |
| 6 | C T G A C T G A G A T C T C A G A G T C A T C G G C T A G T A C A C G T T A G C | 1e-34 | -7.860e+01 | 6.94% | 3.85% | 49.5bp (67.2bp) | MBP1::SWI6/MA0330.1/Jaspar(0.766) More Information \| Similar Motifs Found | motif file (matrix) |
| 7 | C G T A A C T G G T A C C T G A A C G T A T C G A C T G A C G T A G T C A C G T A G T C A G T C | 1e-32 | -7.445e+01 | 0.38% | 0.01% | 31.7bp (48.0bp) | Tv\_0259(RRM)/Trichomonas\_vaginalis-RNCMPT00259-PBM/HughesRNA(0.650) More Information \| Similar Motifs Found | motif file (matrix) |
| 8 | A C G T C T A G G C T A C T G A A C T G C G T A A G T C A C G T C G T A A T C G A T C G A G T C | 1e-28 | -6.611e+01 | 0.47% | 0.03% | 43.5bp (47.4bp) | GATA1/MA0035.5/Jaspar(0.626) More Information \| Similar Motifs Found | motif file (matrix) |
| 9 | A C T G C G T A A G T C A G T C A C G T A C G T G T C A C A T G A G T C A G T C C G A T A T C G | 1e-28 | -6.545e+01 | 0.43% | 0.02% | 43.5bp (39.6bp) | Ppara/MA2338.1/Jaspar(0.739) More Information \| Similar Motifs Found | motif file (matrix) |
| 10 | A C G T A C G T A G T C A G T C C G T A C G T A C G T A A G T C A T C G A C T G C G T A C G T A | 1e-28 | -6.501e+01 | 0.24% | 0.00% | 46.2bp (1.1bp) | insv/MA2315.1/Jaspar(0.735) More Information \| Similar Motifs Found | motif file (matrix) |
| 11 | A C T G A T G C A T C G T A G C C G T A G A C T C G T A G T A C A T C G G A C T | 1e-26 | -6.139e+01 | 2.21% | 0.82% | 51.1bp (64.8bp) | ewg/MA2309.1/Jaspar(0.816) More Information \| Similar Motifs Found | motif file (matrix) |
| 12 | A C G T G A C T G C T A G A C T A G T C A T C G A T G C A C T G A T G C A T C G | 1e-25 | -5.985e+01 | 6.33% | 3.72% | 52.3bp (68.7bp) | LARK(RRM,Znf)/Drosophila\_melanogaster-RNCMPT00035-PBM/HughesRNA(0.899) More Information \| Similar Motifs Found | motif file (matrix) |
| 13 | G A T C T A G C G C A T A C G T T C G A G A C T A G T C C A G T | 1e-25 | -5.786e+01 | 17.90% | 13.51% | 57.1bp (61.6bp) | GATA5/MA0766.3/Jaspar(0.945) More Information \| Similar Motifs Found | motif file (matrix) |
| 14 | A G T C C G T A A G T C C T G A C A T G G A T C A G T C A C T G T C A G G A T C C T G A C A T G | 1e-23 | -5.396e+01 | 0.26% | 0.01% | 25.4bp (47.9bp) | PK24580.1/MA2088.1/Jaspar(0.746) More Information \| Similar Motifs Found | motif file (matrix) |
| 15 | G T A C C G T A G A T C C G T A C A T G T C G A A C G T A C G T G T C A T A C G C G A T C T A G | 1e-22 | -5.140e+01 | 0.63% | 0.09% | 57.8bp (64.9bp) | GATA1/MA0035.5/Jaspar(0.715) More Information \| Similar Motifs Found | motif file (matrix) |
| 16 | T C A G G C A T C T A G G A C T C T A G T C G A C G T A A G T C | 1e-21 | -5.034e+01 | 25.06% | 20.33% | 54.4bp (65.9bp) | Tbx6/MA1567.3/Jaspar(0.755) More Information \| Similar Motifs Found | motif file (matrix) |
| 17 | A T G C C G A T T C G A A C G T C A T G T C G A A T G C C A G T T A G C C T G A | 1e-21 | -4.842e+01 | 18.47% | 14.39% | 53.6bp (61.6bp) | jun-1/MA2135.1/Jaspar(0.891) More Information \| Similar Motifs Found | motif file (matrix) |
| 18 | T A G C A T C G A C G T T C G A T A G C C A T G A C G T C G T A | 1e-20 | -4.795e+01 | 18.41% | 14.35% | 56.0bp (63.3bp) | TFLG2-Zm00001d042777/MA1835.2/Jaspar(0.848) More Information \| Similar Motifs Found | motif file (matrix) |
| 19 | C G A T A G T C C T A G T A C G A G C T G A T C C A T G A T C G | 1e-20 | -4.624e+01 | 5.26% | 3.16% | 51.7bp (66.1bp) | YLL054C/MA0429.1/Jaspar(0.720) More Information \| Similar Motifs Found | motif file (matrix) |
| 20 | A G T C A G T C A C T G A G T C A C T G C G T A A C G T T A C G G A C T A G T C C T A G C G A T | 1e-18 | -4.219e+01 | 0.21% | 0.01% | 57.4bp (63.7bp) | AT5G22990(C2H2)/col-AT5G22990-DAP-Seq(GSE60143)/Homer(0.678) More Information \| Similar Motifs Found | motif file (matrix) |
| 21 | A C T G C T G A A G T C C G T A C G T A C T G A A C G T A C G T C T G A A C G T A G T C A C G T | 1e-18 | -4.160e+01 | 0.19% | 0.01% | 46.0bp (14.6bp) | RVE4/MA1187.2/Jaspar(0.726) More Information \| Similar Motifs Found | motif file (matrix) |
| 22 | C T A G A T G C C G T A T A C G G C T A T A G C A G C T T A G C | 1e-16 | -3.728e+01 | 4.46% | 2.73% | 52.3bp (63.5bp) | Smad3(MAD)/NPC-Smad3-ChIP-Seq(GSE36673)/Homer(0.740) More Information \| Similar Motifs Found | motif file (matrix) |
| 23 | C G T A A C G T A C T G A G T C A C G T A C T G A G T C A G T C | 1e-15 | -3.662e+01 | 0.88% | 0.25% | 50.2bp (57.6bp) | ZNF549/MA1728.2/Jaspar(0.876) More Information \| Similar Motifs Found | motif file (matrix) |
| 24 | T A G C A G T C C G T A A C T G A G C T T A G C A C T G A G C T A C T G T A C G A C G T G C A T | 1e-15 | -3.608e+01 | 0.26% | 0.02% | 50.9bp (33.1bp) | Npas4(bHLH)/Neuron-Npas4-ChIP-Seq(GSE127793)/Homer(0.643) More Information \| Similar Motifs Found | motif file (matrix) |
| 25 | C G T A A C T G A T C G A C T G G T C A A C T G A G T C G A C T T A G C A G T C A C T G A G T C | 1e-14 | -3.438e+01 | 0.36% | 0.04% | 43.4bp (62.5bp) | POL013.1\_MED-1/Jaspar(0.683) More Information \| Similar Motifs Found | motif file (matrix) |
| 26 | C G T A A G T C C G T A C T A G A C T G A C T G C T G A A T G C C G T A C G T A A G C T C T G A | 1e-13 | -3.075e+01 | 0.18% | 0.01% | 48.3bp (39.4bp) | PB0173.1\_Sox21\_2/Jaspar(0.657) More Information \| Similar Motifs Found | motif file (matrix) |
| 27 | A G T C A C T G A C G T A C T G C G T A A C G T A C T G C G T A A C T G A G T C | 1e-12 | -2.957e+01 | 0.21% | 0.01% | 49.0bp (62.4bp) | TOD6?/SacCer-Promoters/Homer(0.768) More Information \| Similar Motifs Found | motif file (matrix) |
| 28 \* | G T C A A C T G A T G C G A T C T A G C C G T A G C T A G T A C A C T G G T A C C G T A T A G C | 1e-10 | -2.474e+01 | 0.14% | 0.01% | 53.1bp (36.0bp) | ZNF449/MA1656.2/Jaspar(0.754) More Information \| Similar Motifs Found | motif file (matrix) |
| 29 \* | A G T C A G C T A G T C G T A C A G T C G T A C A T C G A T G C | 1e-8 | -2.041e+01 | 3.32% | 2.21% | 48.2bp (54.6bp) | rst2/MA1431.2/Jaspar(0.842) More Information \| Similar Motifs Found | motif file (matrix) |
| 30 \* | A G T C C T G A A G T C A C G T A C G T A C T G A C G T A C G T | 1e-8 | -1.860e+01 | 1.99% | 1.19% | 54.5bp (66.2bp) | CDF3/MA0974.3/Jaspar(0.786) More Information \| Similar Motifs Found | motif file (matrix) |
| 31 \* | A C G T T C G A A C T G A C T G A G C T A C G T A C G T C T G A A C T G A T G C | 1e-7 | -1.712e+01 | 0.38% | 0.11% | 51.4bp (65.9bp) | SRS7(SRS)/colamp-SRS7-DAP-Seq(GSE60143)/Homer(0.823) More Information \| Similar Motifs Found | motif file (matrix) |
| 32 \* | C G T A C G T A C G T A C G T A C G T A C G T A C G T A C G T A C G T A C G T A C G T A C G T A | 1e-4 | -1.065e+01 | 1.91% | 1.32% | 42.1bp (58.8bp) | SeqBias: polyA-repeat(0.913) More Information \| Similar Motifs Found | motif file (matrix) |
| 33 \* | A G T C G T A C A G T C G T C A A G T C A G T C A G T C A G T C A G T C A G T C A G T C A T G C | 1e-3 | -7.798e+00 | 12.29% | 11.03% | 53.6bp (56.3bp) | PB0097.1\_Zfp281\_1/Jaspar(0.839) More Information \| Similar Motifs Found | motif file (matrix) |
