## Supplementary_Information for "Genome-wide mapping of the *Galleria mellonella* larvae transcription start sites during fungal infection and treatment": motif1.similar.html

### Information for motif1

A
C
G
T
A
G
T
C
A
C
G
T
C
A
T
G
C
T
G
A
C
A
T
G
A
C
T
G
A
G
T
C
G
T
A
C
C
G
T
A
G
C
T
A
A
G
T
C
  
Reverse Opposite:  

T
A
C
G
C
A
G
T
G
A
C
T
A
C
T
G
C
T
A
G
T
A
G
C
G
T
A
C
A
G
C
T
G
A
T
C
G
T
C
A
C
T
A
G
C
G
T
A
  

|  |  |
| --- | --- |
| p-value: | 1e-63 |
| log p-value: | -1.462e+02 |
| Information Content per bp: | 1.832 |
| Number of Target Sequences with motif | 44.0 |
| Percentage of Target Sequences with motif | 0.61% |
| Number of Background Sequences with motif | 8.8 |
| Percentage of Background Sequences with motif | 0.01% |
| Average Position of motif in Targets | 101.7 +/- 6.0bp |
| Average Position of motif in Background | 108.0 +/- 49.8bp |
| Strand Bias (log2 ratio + to - strand density) | -0.8 |
| Multiplicity (# of sites on avg that occur together) | 1.00 |
| Motif File: | file (matrix) reverse opposite |

#### Similar de novo motifs found

|  |  |  |  |  |  |  |  |
| --- | --- | --- | --- | --- | --- | --- | --- |
| Rank | Match Score | Redundant Motif | P-value | log P-value | % of Targets | % of Background | Motif file |
| 1 | 0.906 | T C A G G C A T A G C T A T C G A C T G T A G C G A T C C G A T T G A C C G T A | 1e-33 | -76.302216 | 1.92% | 0.56% | motif file (matrix) |
| 2 | 0.804 | A T C G A G C T A C G T A C T G A C T G A G T C A G T C A C G T | 1e-23 | -54.389264 | 2.31% | 0.94% | motif file (matrix) |
