## Supplementary_Information for "Genome-wide mapping of the *Galleria mellonella* larvae transcription start sites during fungal infection and treatment": motif2.similar.html

### Information for motif2

A
G
C
T
A
T
G
C
C
G
A
T
A
C
G
T
A
G
T
C
A
G
T
C
A
C
T
G
C
G
T
A
A
C
G
T
A
T
G
C
A
C
G
T
A
T
C
G
  
Reverse Opposite:  

A
T
G
C
C
G
T
A
A
T
C
G
C
G
T
A
A
C
G
T
A
G
T
C
A
C
T
G
C
T
A
G
C
G
T
A
G
C
T
A
T
A
C
G
T
C
G
A
  

|  |  |
| --- | --- |
| p-value: | 1e-51 |
| log p-value: | -1.197e+02 |
| Information Content per bp: | 1.859 |
| Number of Target Sequences with motif | 28.0 |
| Percentage of Target Sequences with motif | 0.39% |
| Number of Background Sequences with motif | 2.5 |
| Percentage of Background Sequences with motif | 0.00% |
| Average Position of motif in Targets | 100.2 +/- 22.6bp |
| Average Position of motif in Background | 106.5 +/- 39.3bp |
| Strand Bias (log2 ratio + to - strand density) | 0.0 |
| Multiplicity (# of sites on avg that occur together) | 1.00 |
| Motif File: | file (matrix) reverse opposite |

#### Similar de novo motifs found

|  |  |  |  |  |  |  |  |
| --- | --- | --- | --- | --- | --- | --- | --- |
| Rank | Match Score | Redundant Motif | P-value | log P-value | % of Targets | % of Background | Motif file |
| 1 | 0.976 | A G C T A C G T A G T C A G T C A C T G C G T A A C G T A G T C A C G T A C T G | 1e-36 | -84.446893 | 0.50% | 0.02% | motif file (matrix) |
| 2 | 0.878 | G T A C A G T C A C T G C G T A A C G T A G T C A C G T A C T G | 1e-32 | -74.875500 | 1.60% | 0.41% | motif file (matrix) |
| 3 | 0.800 | A C G T A G T C A G T C A C T G C G T A A G T C A G T C A C G T | 1e-30 | -70.069662 | 1.32% | 0.30% | motif file (matrix) |
| 4 | 0.775 | C G T A T A G C T C G A T C A G G C T A A G C T T A G C C A T G C A T G C T A G | 1e-25 | -57.715425 | 5.92% | 3.45% | motif file (matrix) |
| 5 | 0.718 | A G T C A C G T A G T C T A C G A T G C A T C G C G T A A C G T A G T C A C G T A C T G C G T A | 1e-20 | -46.673363 | 0.18% | 0.00% | motif file (matrix) |
| 6 | 0.717 | T G A C G C T A C A T G G C T A G A C T G T A C C T G A G T A C C G T A G T C A | 1e-19 | -45.426752 | 13.85% | 10.39% | motif file (matrix) |
