## Supplementary_Information for "Genome-wide mapping of the *Galleria mellonella* larvae transcription start sites during fungal infection and treatment": motif3.similar.html

### Information for motif3

T
A
C
G
A
C
T
G
A
C
G
T
A
T
G
C
G
T
C
A
T
C
A
G
A
C
G
T
T
C
A
G
  
Reverse Opposite:  

A
G
T
C
G
T
C
A
A
G
T
C
A
C
G
T
T
A
C
G
T
G
C
A
T
G
A
C
A
T
G
C
  

|  |  |
| --- | --- |
| p-value: | 1e-40 |
| log p-value: | -9.353e+01 |
| Information Content per bp: | 1.796 |
| Number of Target Sequences with motif | 2849.0 |
| Percentage of Target Sequences with motif | 39.62% |
| Number of Background Sequences with motif | 29161.4 |
| Percentage of Background Sequences with motif | 32.10% |
| Average Position of motif in Targets | 100.7 +/- 51.9bp |
| Average Position of motif in Background | 100.3 +/- 64.0bp |
| Strand Bias (log2 ratio + to - strand density) | -0.0 |
| Multiplicity (# of sites on avg that occur together) | 1.32 |
| Motif File: | file (matrix) reverse opposite |

#### Similar de novo motifs found

|  |  |  |  |  |  |  |  |
| --- | --- | --- | --- | --- | --- | --- | --- |
| Rank | Match Score | Redundant Motif | P-value | log P-value | % of Targets | % of Background | Motif file |
| 1 | 0.797 | G C A T T C A G C G A T G T A C C G T A T A C G G A C T T C A G G C T A A G T C | 1e-29 | -68.034322 | 10.21% | 6.62% | motif file (matrix) |
| 2 | 0.792 | G C T A G C A T C T A G G C T A G T A C C G T A G T A C G C A T C T A G C G T A G A T C C G T A | 1e-22 | -51.802393 | 22.77% | 18.15% | motif file (matrix) |
| 3 | 0.681 | G T A C C G T A T C G A A G T C T G C A T A G C C G A T C T A G C T G A A G C T | 1e-20 | -46.081815 | 13.52% | 10.07% | motif file (matrix) |
| 4 | 0.722 | A T C G T G A C C T A G T C G A C A G T T A C G T C G A A T C G G A C T C A T G | 1e-15 | -36.492829 | 6.30% | 4.22% | motif file (matrix) |
| 5 | 0.628 | A C G T A C G T A C G T A G T C C G T A A C T G A C G T A C T G A G T C A C T G | 1e-8 | -20.254724 | 0.14% | 0.01% | motif file (matrix) |
