## Supplementary_Information for "Genome-wide mapping of the *Galleria mellonella* larvae transcription start sites during fungal infection and treatment": motif4.similar.html

### Information for motif4

T
C
A
G
T
G
A
C
A
T
G
C
T
C
G
A
C
T
G
A
A
G
C
T
C
T
A
G
G
C
A
T
G
A
T
C
C
T
G
A
  
Reverse Opposite:  

A
G
C
T
C
T
A
G
C
G
T
A
G
A
T
C
T
C
G
A
G
A
C
T
A
G
C
T
T
A
C
G
A
C
T
G
A
G
T
C
  

|  |  |
| --- | --- |
| p-value: | 1e-38 |
| log p-value: | -8.806e+01 |
| Information Content per bp: | 1.579 |
| Number of Target Sequences with motif | 1580.0 |
| Percentage of Target Sequences with motif | 21.97% |
| Number of Background Sequences with motif | 14613.3 |
| Percentage of Background Sequences with motif | 16.09% |
| Average Position of motif in Targets | 100.7 +/- 55.3bp |
| Average Position of motif in Background | 100.6 +/- 63.0bp |
| Strand Bias (log2 ratio + to - strand density) | 0.1 |
| Multiplicity (# of sites on avg that occur together) | 1.10 |
| Motif File: | file (matrix) reverse opposite |

#### Similar de novo motifs found

|  |  |  |  |  |  |  |  |
| --- | --- | --- | --- | --- | --- | --- | --- |
| Rank | Match Score | Redundant Motif | P-value | log P-value | % of Targets | % of Background | Motif file |
| 1 | 0.701 | A G C T T C A G C G T A G A T C T G A C A G C T A C G T T C A G | 1e-29 | -67.670362 | 28.66% | 22.89% | motif file (matrix) |
