## Supplementary_Information for "Genome-wide mapping of the *Galleria mellonella* larvae transcription start sites during fungal infection and treatment": motif5.similar.html

### Information for motif5

A
T
G
C
T
A
G
C
A
T
G
C
C
T
A
G
A
C
G
T
A
G
T
C
G
T
C
A
A
C
T
G
C
G
A
T
G
T
A
C
  
Reverse Opposite:  

C
A
T
G
C
G
T
A
T
G
A
C
C
A
G
T
A
C
T
G
T
G
C
A
G
A
T
C
T
A
C
G
A
T
C
G
A
T
C
G
  

|  |  |
| --- | --- |
| p-value: | 1e-35 |
| log p-value: | -8.223e+01 |
| Information Content per bp: | 1.575 |
| Number of Target Sequences with motif | 1610.0 |
| Percentage of Target Sequences with motif | 22.39% |
| Number of Background Sequences with motif | 15121.8 |
| Percentage of Background Sequences with motif | 16.64% |
| Average Position of motif in Targets | 102.1 +/- 51.1bp |
| Average Position of motif in Background | 98.9 +/- 63.1bp |
| Strand Bias (log2 ratio + to - strand density) | 0.0 |
| Multiplicity (# of sites on avg that occur together) | 1.17 |
| Motif File: | file (matrix) reverse opposite |

#### Similar de novo motifs found

|  |  |  |  |  |  |  |  |
| --- | --- | --- | --- | --- | --- | --- | --- |
| Rank | Match Score | Redundant Motif | P-value | log P-value | % of Targets | % of Background | Motif file |
| 1 | 0.622 | T G C A A C G T A C T G T C G A A G T C C T A G G A C T G A T C | 1e-20 | -47.538414 | 12.74% | 9.34% | motif file (matrix) |
| 2 | 0.654 | C T A G C G T A A T G C A G C T A C T G C T A G T G C A C A T G | 1e-13 | -31.619483 | 7.72% | 5.56% | motif file (matrix) |
