## Supplementary_Information for "Genome-wide mapping of the *Galleria mellonella* larvae transcription start sites during fungal infection and treatment": motif6.similar.html

### Information for motif6

C
T
G
A
C
T
G
A
G
A
T
C
T
C
A
G
A
G
T
C
A
T
C
G
G
C
T
A
G
T
A
C
A
C
G
T
T
A
G
C
  
Reverse Opposite:  

A
T
C
G
G
T
C
A
C
A
T
G
C
G
A
T
T
A
G
C
T
C
A
G
A
G
T
C
C
T
A
G
G
A
C
T
G
A
C
T
  

|  |  |
| --- | --- |
| p-value: | 1e-34 |
| log p-value: | -7.860e+01 |
| Information Content per bp: | 1.648 |
| Number of Target Sequences with motif | 499.0 |
| Percentage of Target Sequences with motif | 6.94% |
| Number of Background Sequences with motif | 3499.9 |
| Percentage of Background Sequences with motif | 3.85% |
| Average Position of motif in Targets | 100.7 +/- 49.5bp |
| Average Position of motif in Background | 100.2 +/- 67.2bp |
| Strand Bias (log2 ratio + to - strand density) | -0.1 |
| Multiplicity (# of sites on avg that occur together) | 1.04 |
| Motif File: | file (matrix) reverse opposite |

#### Similar de novo motifs found

|  |  |  |  |  |  |  |  |
| --- | --- | --- | --- | --- | --- | --- | --- |
| Rank | Match Score | Redundant Motif | P-value | log P-value | % of Targets | % of Background | Motif file |
| 1 | 0.805 | A T C G G T C A A C T G C G T A T G A C A C T G A G T C A C T G | 1e-26 | -60.693795 | 20.08% | 15.35% | motif file (matrix) |
| 2 | 0.765 | C A G T T A G C C G T A C A T G C G A T T A G C C T A G G T C A T C A G G C A T G A C T C T A G | 1e-20 | -48.227369 | 1.57% | 0.55% | motif file (matrix) |
| 3 | 0.677 | C T A G G T A C C A T G G T A C A T C G T G C A A G T C C A G T T G A C C A G T A G C T T A C G | 1e-16 | -39.138344 | 0.89% | 0.25% | motif file (matrix) |
| 4 | 0.603 | G T A C A C G T A G T C A C T G A G T C A C T G C G T A A C G T A C T G A G C T | 1e-10 | -24.330747 | 0.35% | 0.06% | motif file (matrix) |
