## Supplementary_Information for "Genome-wide mapping of the *Galleria mellonella* larvae transcription start sites during fungal infection and treatment": motif7.similar.html

### Information for motif7

C
G
T
A
A
C
T
G
G
T
A
C
C
T
G
A
A
C
G
T
A
T
C
G
A
C
T
G
A
C
G
T
A
G
T
C
A
C
G
T
A
G
T
C
A
G
T
C
  
Reverse Opposite:  

A
C
T
G
A
C
T
G
C
G
T
A
A
C
T
G
C
G
T
A
A
G
T
C
A
T
G
C
C
G
T
A
A
G
C
T
A
C
T
G
A
G
T
C
A
C
G
T
  

|  |  |
| --- | --- |
| p-value: | 1e-32 |
| log p-value: | -7.445e+01 |
| Information Content per bp: | 1.971 |
| Number of Target Sequences with motif | 27.0 |
| Percentage of Target Sequences with motif | 0.38% |
| Number of Background Sequences with motif | 9.3 |
| Percentage of Background Sequences with motif | 0.01% |
| Average Position of motif in Targets | 126.1 +/- 31.7bp |
| Average Position of motif in Background | 122.7 +/- 48.0bp |
| Strand Bias (log2 ratio + to - strand density) | -0.1 |
| Multiplicity (# of sites on avg that occur together) | 1.00 |
| Motif File: | file (matrix) reverse opposite |

#### Similar de novo motifs found

|  |  |  |  |  |  |  |  |
| --- | --- | --- | --- | --- | --- | --- | --- |
| Rank | Match Score | Redundant Motif | P-value | log P-value | % of Targets | % of Background | Motif file |
| 1 | 0.818 | A G T C C T G A A C T G A G T C C G T A A C G T A C T G A C T G A C G T A G T C | 1e-24 | -56.906954 | 0.40% | 0.02% | motif file (matrix) |
