## Supplementary_Information for "Genome-wide mapping of the *Galleria mellonella* larvae transcription start sites during fungal infection and treatment": motif8.similar.html

### Information for motif8

A
C
G
T
C
T
A
G
G
C
T
A
C
T
G
A
A
C
T
G
C
G
T
A
A
G
T
C
A
C
G
T
C
G
T
A
A
T
C
G
A
T
C
G
A
G
T
C
  
Reverse Opposite:  

A
C
T
G
A
T
G
C
A
T
G
C
G
C
A
T
C
G
T
A
C
T
A
G
A
C
G
T
A
G
T
C
A
G
C
T
C
G
A
T
A
G
T
C
C
G
T
A
  

|  |  |
| --- | --- |
| p-value: | 1e-28 |
| log p-value: | -6.611e+01 |
| Information Content per bp: | 1.821 |
| Number of Target Sequences with motif | 34.0 |
| Percentage of Target Sequences with motif | 0.47% |
| Number of Background Sequences with motif | 26.1 |
| Percentage of Background Sequences with motif | 0.03% |
| Average Position of motif in Targets | 105.8 +/- 43.5bp |
| Average Position of motif in Background | 119.3 +/- 47.4bp |
| Strand Bias (log2 ratio + to - strand density) | 0.5 |
| Multiplicity (# of sites on avg that occur together) | 1.00 |
| Motif File: | file (matrix) reverse opposite |

#### Similar de novo motifs found

|  |  |  |  |  |  |  |  |
| --- | --- | --- | --- | --- | --- | --- | --- |
| Rank | Match Score | Redundant Motif | P-value | log P-value | % of Targets | % of Background | Motif file |
| 1 | 0.606 | A T C G G T A C C G T A T C A G G T C A A G T C A G C T G C T A C T A G A C T G | 1e-17 | -39.950774 | 0.86% | 0.23% | motif file (matrix) |
