## Supplementary_Information for "Genome-wide mapping of the *Galleria mellonella* larvae transcription start sites during fungal infection and treatment": motif9.similar.html

### Information for motif9

A
C
T
G
C
G
T
A
A
G
T
C
A
G
T
C
A
C
G
T
A
C
G
T
G
T
C
A
C
A
T
G
A
G
T
C
A
G
T
C
C
G
A
T
A
T
C
G
  
Reverse Opposite:  

A
T
G
C
C
G
T
A
A
C
T
G
A
C
T
G
G
A
T
C
A
C
G
T
C
G
T
A
C
G
T
A
C
T
A
G
A
C
T
G
A
C
G
T
A
G
T
C
  

|  |  |
| --- | --- |
| p-value: | 1e-28 |
| log p-value: | -6.545e+01 |
| Information Content per bp: | 1.844 |
| Number of Target Sequences with motif | 31.0 |
| Percentage of Target Sequences with motif | 0.43% |
| Number of Background Sequences with motif | 20.0 |
| Percentage of Background Sequences with motif | 0.02% |
| Average Position of motif in Targets | 119.9 +/- 43.5bp |
| Average Position of motif in Background | 107.8 +/- 39.6bp |
| Strand Bias (log2 ratio + to - strand density) | 0.0 |
| Multiplicity (# of sites on avg that occur together) | 1.03 |
| Motif File: | file (matrix) reverse opposite |

#### Similar de novo motifs found

|  |  |  |  |  |  |  |  |
| --- | --- | --- | --- | --- | --- | --- | --- |
| Rank | Match Score | Redundant Motif | P-value | log P-value | % of Targets | % of Background | Motif file |
| 1 | 0.682 | A G T C A T G C A C T G C G T A A G T C A G T C A C G T A C G T C G T A A C T G | 1e-27 | -62.466389 | 0.42% | 0.02% | motif file (matrix) |
