## Supplementary_Information for "Genome-wide mapping of the *Galleria mellonella* larvae transcription start sites during fungal infection and treatment": motif10.similar.html

### Information for motif10

A
C
G
T
A
C
G
T
A
G
T
C
A
G
T
C
C
G
T
A
C
G
T
A
C
G
T
A
A
G
T
C
A
T
C
G
A
C
T
G
C
G
T
A
C
G
T
A
  
Reverse Opposite:  

A
C
G
T
A
C
G
T
A
G
T
C
A
T
G
C
A
C
T
G
A
C
G
T
A
C
G
T
A
C
G
T
A
C
T
G
A
C
T
G
C
G
T
A
C
G
T
A
  

|  |  |
| --- | --- |
| p-value: | 1e-28 |
| log p-value: | -6.501e+01 |
| Information Content per bp: | 1.982 |
| Number of Target Sequences with motif | 17.0 |
| Percentage of Target Sequences with motif | 0.24% |
| Number of Background Sequences with motif | 2.3 |
| Percentage of Background Sequences with motif | 0.00% |
| Average Position of motif in Targets | 104.3 +/- 46.2bp |
| Average Position of motif in Background | 110.0 +/- 1.1bp |
| Strand Bias (log2 ratio + to - strand density) | -0.2 |
| Multiplicity (# of sites on avg that occur together) | 1.00 |
| Motif File: | file (matrix) reverse opposite |

#### Similar de novo motifs found

|  |  |  |  |  |  |  |  |
| --- | --- | --- | --- | --- | --- | --- | --- |
| Rank | Match Score | Redundant Motif | P-value | log P-value | % of Targets | % of Background | Motif file |
| 1 | 0.912 | A C G T A G T C A G T C C G T A C G T A C G T A A G T C A C T G A C T G C G T A | 1e-16 | -36.875764 | 0.25% | 0.02% | motif file (matrix) |
