## Supplementary_Information for "Genome-wide mapping of the *Galleria mellonella* larvae transcription start sites during fungal infection and treatment": motif11.similar.html

### Information for motif11

A
C
T
G
A
T
G
C
A
T
C
G
T
A
G
C
C
G
T
A
G
A
C
T
C
G
T
A
G
T
A
C
A
T
C
G
G
A
C
T
  
Reverse Opposite:  

C
T
G
A
A
T
G
C
C
A
T
G
G
C
A
T
C
T
G
A
C
G
A
T
A
T
C
G
T
A
G
C
A
T
C
G
G
T
A
C
  

|  |  |
| --- | --- |
| p-value: | 1e-26 |
| log p-value: | -6.139e+01 |
| Information Content per bp: | 1.688 |
| Number of Target Sequences with motif | 159.0 |
| Percentage of Target Sequences with motif | 2.21% |
| Number of Background Sequences with motif | 745.9 |
| Percentage of Background Sequences with motif | 0.82% |
| Average Position of motif in Targets | 103.6 +/- 51.1bp |
| Average Position of motif in Background | 96.7 +/- 64.8bp |
| Strand Bias (log2 ratio + to - strand density) | 0.1 |
| Multiplicity (# of sites on avg that occur together) | 1.03 |
| Motif File: | file (matrix) reverse opposite |

#### Similar de novo motifs found

|  |  |  |  |  |  |  |  |
| --- | --- | --- | --- | --- | --- | --- | --- |
| Rank | Match Score | Redundant Motif | P-value | log P-value | % of Targets | % of Background | Motif file |
| 1 | 0.728 | G T C A A G T C G T C A G T A C A C T G C G T A C G T A C G A T A T G C C T A G A C T G G T A C | 1e-24 | -57.383822 | 0.50% | 0.05% | motif file (matrix) |
| 2 | 0.662 | C A G T A T G C A C T G A T G C C T G A G C A T T A G C A G T C | 1e-13 | -30.873951 | 2.89% | 1.65% | motif file (matrix) |
| 3 | 0.803 | G A T C C T A G A G C T C G T A A G C T C A T G G A T C C T A G G A T C T C G A G T A C C G A T | 1e-11 | -26.577064 | 2.24% | 1.24% | motif file (matrix) |
