## Supplementary_Information for "Genome-wide mapping of the *Galleria mellonella* larvae transcription start sites during fungal infection and treatment": motif12.similar.html

### Information for motif12

A
C
G
T
G
A
C
T
G
C
T
A
G
A
C
T
A
G
T
C
A
T
C
G
A
T
G
C
A
C
T
G
A
T
G
C
A
T
C
G
  
Reverse Opposite:  

T
A
G
C
A
T
C
G
G
T
A
C
T
A
C
G
A
T
G
C
C
T
A
G
C
T
G
A
C
G
A
T
C
T
G
A
T
G
C
A
  

|  |  |
| --- | --- |
| p-value: | 1e-25 |
| log p-value: | -5.985e+01 |
| Information Content per bp: | 1.686 |
| Number of Target Sequences with motif | 455.0 |
| Percentage of Target Sequences with motif | 6.33% |
| Number of Background Sequences with motif | 3380.4 |
| Percentage of Background Sequences with motif | 3.72% |
| Average Position of motif in Targets | 102.4 +/- 52.3bp |
| Average Position of motif in Background | 101.1 +/- 68.7bp |
| Strand Bias (log2 ratio + to - strand density) | 0.0 |
| Multiplicity (# of sites on avg that occur together) | 1.08 |
| Motif File: | file (matrix) reverse opposite |

#### Similar de novo motifs found

|  |  |  |  |  |  |  |  |
| --- | --- | --- | --- | --- | --- | --- | --- |
| Rank | Match Score | Redundant Motif | P-value | log P-value | % of Targets | % of Background | Motif file |
| 1 | 0.822 | A C G T A C G T G C A T A C G T A T G C T A G C G A T C C A T G T G A C A C T G A G C T C G A T | 1e-21 | -50.498774 | 2.02% | 0.80% | motif file (matrix) |
| 2 | 0.703 | T A G C T A C G G T A C C A T G C T A G G C A T T G C A T G C A | 1e-20 | -47.023494 | 21.77% | 17.44% | motif file (matrix) |
| 3 | 0.603 | A G C T A T G C A T G C A T G C T A C G G T A C A T G C G T A C | 1e-9 | -22.376761 | 7.08% | 5.34% | motif file (matrix) |
