## Supplementary_Information for "Genome-wide mapping of the *Galleria mellonella* larvae transcription start sites during fungal infection and treatment": motif13.similar.html

### Information for motif13

G
A
T
C
T
A
G
C
G
C
A
T
A
C
G
T
T
C
G
A
G
A
C
T
A
G
T
C
C
A
G
T
  
Reverse Opposite:  

G
T
C
A
A
C
T
G
C
T
G
A
A
G
C
T
G
T
C
A
C
G
T
A
A
T
C
G
C
T
A
G
  

|  |  |
| --- | --- |
| p-value: | 1e-25 |
| log p-value: | -5.786e+01 |
| Information Content per bp: | 1.651 |
| Number of Target Sequences with motif | 1287.0 |
| Percentage of Target Sequences with motif | 17.90% |
| Number of Background Sequences with motif | 12271.9 |
| Percentage of Background Sequences with motif | 13.51% |
| Average Position of motif in Targets | 97.5 +/- 57.1bp |
| Average Position of motif in Background | 100.1 +/- 61.6bp |
| Strand Bias (log2 ratio + to - strand density) | 0.1 |
| Multiplicity (# of sites on avg that occur together) | 1.11 |
| Motif File: | file (matrix) reverse opposite |

#### Similar de novo motifs found

|  |  |  |  |  |  |  |  |
| --- | --- | --- | --- | --- | --- | --- | --- |
| Rank | Match Score | Redundant Motif | P-value | log P-value | % of Targets | % of Background | Motif file |
| 1 | 0.693 | C G A T G A C T T C G A A G C T T G A C C A G T C T G A A T C G C A T G C A G T | 1e-20 | -46.672342 | 29.37% | 24.54% | motif file (matrix) |
| 2 | 0.643 | T G A C A T C G C T G A A T G C C G A T A C T G G T C A A G C T C G T A T C G A | 1e-16 | -37.636743 | 1.60% | 0.66% | motif file (matrix) |
| 3 | 0.631 | C T A G G A C T G A T C G C T A C G A T T G A C G C A T C T A G | 1e-16 | -37.463516 | 7.45% | 5.15% | motif file (matrix) |
