## Supplementary_Information for "Genome-wide mapping of the *Galleria mellonella* larvae transcription start sites during fungal infection and treatment": motif14.similar.html

### Information for motif14

A
G
T
C
C
G
T
A
A
G
T
C
C
T
G
A
C
A
T
G
G
A
T
C
A
G
T
C
A
C
T
G
T
C
A
G
G
A
T
C
C
T
G
A
C
A
T
G
  
Reverse Opposite:  

G
T
A
C
G
A
C
T
C
T
A
G
A
G
T
C
T
G
A
C
C
T
A
G
C
T
A
G
G
T
A
C
G
A
C
T
T
C
A
G
G
C
A
T
C
T
A
G
  

|  |  |
| --- | --- |
| p-value: | 1e-23 |
| log p-value: | -5.396e+01 |
| Information Content per bp: | 1.625 |
| Number of Target Sequences with motif | 19.0 |
| Percentage of Target Sequences with motif | 0.26% |
| Number of Background Sequences with motif | 6.1 |
| Percentage of Background Sequences with motif | 0.01% |
| Average Position of motif in Targets | 109.1 +/- 25.4bp |
| Average Position of motif in Background | 117.3 +/- 47.9bp |
| Strand Bias (log2 ratio + to - strand density) | -0.7 |
| Multiplicity (# of sites on avg that occur together) | 1.00 |
| Motif File: | file (matrix) reverse opposite |

#### Similar de novo motifs found

|  |  |  |  |  |  |  |  |
| --- | --- | --- | --- | --- | --- | --- | --- |
| Rank | Match Score | Redundant Motif | P-value | log P-value | % of Targets | % of Background | Motif file |
