## Supplementary_Information for "Genome-wide mapping of the *Galleria mellonella* larvae transcription start sites during fungal infection and treatment": motif15.similar.html

### Information for motif15

G
T
A
C
C
G
T
A
G
A
T
C
C
G
T
A
C
A
T
G
T
C
G
A
A
C
G
T
A
C
G
T
G
T
C
A
T
A
C
G
C
G
A
T
C
T
A
G
  
Reverse Opposite:  

A
G
T
C
C
G
T
A
A
T
G
C
A
C
G
T
G
T
C
A
G
T
C
A
A
C
G
T
G
T
A
C
C
G
A
T
C
T
A
G
G
A
C
T
A
C
T
G
  

|  |  |
| --- | --- |
| p-value: | 1e-22 |
| log p-value: | -5.140e+01 |
| Information Content per bp: | 1.748 |
| Number of Target Sequences with motif | 45.0 |
| Percentage of Target Sequences with motif | 0.63% |
| Number of Background Sequences with motif | 82.1 |
| Percentage of Background Sequences with motif | 0.09% |
| Average Position of motif in Targets | 89.6 +/- 57.8bp |
| Average Position of motif in Background | 98.6 +/- 64.9bp |
| Strand Bias (log2 ratio + to - strand density) | 1.2 |
| Multiplicity (# of sites on avg that occur together) | 1.00 |
| Motif File: | file (matrix) reverse opposite |

#### Similar de novo motifs found

|  |  |  |  |  |  |  |  |
| --- | --- | --- | --- | --- | --- | --- | --- |
| Rank | Match Score | Redundant Motif | P-value | log P-value | % of Targets | % of Background | Motif file |
| 1 | 0.761 | T A G C G A T C C G T A A T G C C T G A A C T G C T G A A C G T | 1e-19 | -45.839674 | 3.30% | 1.70% | motif file (matrix) |
