## Supplementary_Information for "Genome-wide mapping of the *Galleria mellonella* larvae transcription start sites during fungal infection and treatment": motif17.similar.html

### Information for motif17

A
T
G
C
C
G
A
T
T
C
G
A
A
C
G
T
C
A
T
G
T
C
G
A
A
T
G
C
C
A
G
T
T
A
G
C
C
T
G
A
  
Reverse Opposite:  

G
A
C
T
A
T
C
G
G
T
C
A
T
A
C
G
A
G
C
T
G
A
T
C
T
G
C
A
A
G
C
T
G
C
T
A
T
A
C
G
  

|  |  |
| --- | --- |
| p-value: | 1e-21 |
| log p-value: | -4.842e+01 |
| Information Content per bp: | 1.532 |
| Number of Target Sequences with motif | 1328.0 |
| Percentage of Target Sequences with motif | 18.47% |
| Number of Background Sequences with motif | 13069.2 |
| Percentage of Background Sequences with motif | 14.39% |
| Average Position of motif in Targets | 100.4 +/- 53.6bp |
| Average Position of motif in Background | 99.6 +/- 61.6bp |
| Strand Bias (log2 ratio + to - strand density) | 0.0 |
| Multiplicity (# of sites on avg that occur together) | 1.13 |
| Motif File: | file (matrix) reverse opposite |

#### Similar de novo motifs found

|  |  |  |  |  |  |  |  |
| --- | --- | --- | --- | --- | --- | --- | --- |
| Rank | Match Score | Redundant Motif | P-value | log P-value | % of Targets | % of Background | Motif file |
| 1 | 0.776 | A G C T A T C G G T C A A T G C A C G T A T G C C T G A C A G T | 1e-20 | -46.949807 | 15.92% | 12.17% | motif file (matrix) |
| 2 | 0.672 | G C T A C T A G A G T C T C A G G C T A G A C T T C A G C G T A T G A C C G A T C T A G C G T A | 1e-19 | -45.627571 | 1.70% | 0.65% | motif file (matrix) |
| 3 | 0.783 | G T C A A C G T A C T G C G T A C G A T A C G T A G T C C G T A | 1e-8 | -20.467512 | 2.29% | 1.39% | motif file (matrix) |
