## Supplementary_Information for "Genome-wide mapping of the *Galleria mellonella* larvae transcription start sites during fungal infection and treatment": motif18.similar.html

### Information for motif18

T
A
G
C
A
T
C
G
A
C
G
T
T
C
G
A
T
A
G
C
C
A
T
G
A
C
G
T
C
G
T
A
  
Reverse Opposite:  

C
G
A
T
C
G
T
A
G
T
A
C
A
T
C
G
A
G
C
T
C
G
T
A
A
T
G
C
A
T
C
G
  

|  |  |
| --- | --- |
| p-value: | 1e-20 |
| log p-value: | -4.795e+01 |
| Information Content per bp: | 1.666 |
| Number of Target Sequences with motif | 1324.0 |
| Percentage of Target Sequences with motif | 18.41% |
| Number of Background Sequences with motif | 13041.1 |
| Percentage of Background Sequences with motif | 14.35% |
| Average Position of motif in Targets | 98.5 +/- 56.0bp |
| Average Position of motif in Background | 100.3 +/- 63.3bp |
| Strand Bias (log2 ratio + to - strand density) | 0.0 |
| Multiplicity (# of sites on avg that occur together) | 1.13 |
| Motif File: | file (matrix) reverse opposite |

#### Similar de novo motifs found

|  |  |  |  |  |  |  |  |
| --- | --- | --- | --- | --- | --- | --- | --- |
| Rank | Match Score | Redundant Motif | P-value | log P-value | % of Targets | % of Background | Motif file |
