## Supplementary_Information for "Genome-wide mapping of the *Galleria mellonella* larvae transcription start sites during fungal infection and treatment": motif24.similar.html

### Information for motif24

T
A
G
C
A
G
T
C
C
G
T
A
A
C
T
G
A
G
C
T
T
A
G
C
A
C
T
G
A
G
C
T
A
C
T
G
T
A
C
G
A
C
G
T
G
C
A
T
  
Reverse Opposite:  

C
G
T
A
T
C
G
A
A
T
G
C
T
G
A
C
C
T
G
A
A
G
T
C
A
T
C
G
C
T
G
A
G
T
A
C
C
G
A
T
A
C
T
G
A
T
C
G
  

|  |  |
| --- | --- |
| p-value: | 1e-15 |
| log p-value: | -3.608e+01 |
| Information Content per bp: | 1.794 |
| Number of Target Sequences with motif | 19.0 |
| Percentage of Target Sequences with motif | 0.26% |
| Number of Background Sequences with motif | 16.8 |
| Percentage of Background Sequences with motif | 0.02% |
| Average Position of motif in Targets | 82.3 +/- 50.9bp |
| Average Position of motif in Background | 115.4 +/- 33.1bp |
| Strand Bias (log2 ratio + to - strand density) | 0.7 |
| Multiplicity (# of sites on avg that occur together) | 1.17 |
| Motif File: | file (matrix) reverse opposite |

#### Similar de novo motifs found

|  |  |  |  |  |  |  |  |
| --- | --- | --- | --- | --- | --- | --- | --- |
| Rank | Match Score | Redundant Motif | P-value | log P-value | % of Targets | % of Background | Motif file |
