## Supplementary_Information for "Genome-wide mapping of the *Galleria mellonella* larvae transcription start sites during fungal infection and treatment": motif27.similar.html

### Information for motif27

A
G
T
C
A
C
T
G
A
C
G
T
A
C
T
G
C
G
T
A
A
C
G
T
A
C
T
G
C
G
T
A
A
C
T
G
A
G
T
C
  
Reverse Opposite:  

A
C
T
G
A
G
T
C
A
C
G
T
G
T
A
C
C
G
T
A
A
C
G
T
A
G
T
C
C
G
T
A
A
G
T
C
A
C
T
G
  

|  |  |
| --- | --- |
| p-value: | 1e-12 |
| log p-value: | -2.957e+01 |
| Information Content per bp: | 1.964 |
| Number of Target Sequences with motif | 15.0 |
| Percentage of Target Sequences with motif | 0.21% |
| Number of Background Sequences with motif | 12.7 |
| Percentage of Background Sequences with motif | 0.01% |
| Average Position of motif in Targets | 72.1 +/- 49.0bp |
| Average Position of motif in Background | 95.9 +/- 62.4bp |
| Strand Bias (log2 ratio + to - strand density) | 1.0 |
| Multiplicity (# of sites on avg that occur together) | 1.00 |
| Motif File: | file (matrix) reverse opposite |

#### Similar de novo motifs found

|  |  |  |  |  |  |  |  |
| --- | --- | --- | --- | --- | --- | --- | --- |
| Rank | Match Score | Redundant Motif | P-value | log P-value | % of Targets | % of Background | Motif file |
