## Supplementary_Information for "Genome-wide mapping of the *Galleria mellonella* larvae transcription start sites during fungal infection and treatment": motif29.similar.html

### Information for motif29

A
G
T
C
A
G
C
T
A
G
T
C
G
T
A
C
A
G
T
C
G
T
A
C
A
T
C
G
A
T
G
C
  
Reverse Opposite:  

A
T
C
G
A
T
G
C
A
C
T
G
A
C
T
G
A
C
T
G
A
C
T
G
C
T
G
A
A
C
T
G
  

|  |  |
| --- | --- |
| p-value: | 1e-8 |
| log p-value: | -2.041e+01 |
| Information Content per bp: | 1.937 |
| Number of Target Sequences with motif | 239.0 |
| Percentage of Target Sequences with motif | 3.32% |
| Number of Background Sequences with motif | 2010.1 |
| Percentage of Background Sequences with motif | 2.21% |
| Average Position of motif in Targets | 99.4 +/- 48.2bp |
| Average Position of motif in Background | 101.1 +/- 54.6bp |
| Strand Bias (log2 ratio + to - strand density) | -0.2 |
| Multiplicity (# of sites on avg that occur together) | 1.06 |
| Motif File: | file (matrix) reverse opposite |

#### Similar de novo motifs found

|  |  |  |  |  |  |  |  |
| --- | --- | --- | --- | --- | --- | --- | --- |
| Rank | Match Score | Redundant Motif | P-value | log P-value | % of Targets | % of Background | Motif file |
| 1 | 0.736 | C A T G T C A G A C T G A T G C C T A G A T C G A C T G A T C G C T A G A C T G | 1e-8 | -19.697197 | 1.22% | 0.61% | motif file (matrix) |
