## Supplementary_Information for "Genome-wide mapping of the *Galleria mellonella* larvae transcription start sites during fungal infection and treatment": motif31.similar.html

### Information for motif31

A
C
G
T
T
C
G
A
A
C
T
G
A
C
T
G
A
G
C
T
A
C
G
T
A
C
G
T
C
T
G
A
A
C
T
G
A
T
G
C
  
Reverse Opposite:  

A
T
C
G
G
T
A
C
G
A
C
T
C
G
T
A
T
G
C
A
C
T
G
A
A
G
T
C
A
G
T
C
A
G
C
T
G
T
C
A
  

|  |  |
| --- | --- |
| p-value: | 1e-7 |
| log p-value: | -1.712e+01 |
| Information Content per bp: | 1.782 |
| Number of Target Sequences with motif | 27.0 |
| Percentage of Target Sequences with motif | 0.38% |
| Number of Background Sequences with motif | 96.0 |
| Percentage of Background Sequences with motif | 0.11% |
| Average Position of motif in Targets | 112.1 +/- 51.4bp |
| Average Position of motif in Background | 115.4 +/- 65.9bp |
| Strand Bias (log2 ratio + to - strand density) | 1.3 |
| Multiplicity (# of sites on avg that occur together) | 1.30 |
| Motif File: | file (matrix) reverse opposite |

#### Similar de novo motifs found

|  |  |  |  |  |  |  |  |
| --- | --- | --- | --- | --- | --- | --- | --- |
| Rank | Match Score | Redundant Motif | P-value | log P-value | % of Targets | % of Background | Motif file |
| 1 | 0.667 | C G T A A G T C A G T C A C G T C G T A C G T A C G T A A G T C | 1e-4 | -10.134842 | 0.79% | 0.45% | motif file (matrix) |
| 2 | 0.736 | A C T G A C T G A C G T A C G T A C G T C G T A A C T G A C T G A C G T A C G T A C G T C G T A | 1e0 | -2.110604 | 72.63% | 72.00% | motif file (matrix) |
