## Supplementary_Information for "Genome-wide mapping of the *Galleria mellonella* larvae transcription start sites during fungal infection and treatment": motif32.similar.html

### Information for motif32

C
G
T
A
C
G
T
A
C
G
T
A
C
G
T
A
C
G
T
A
C
G
T
A
C
G
T
A
C
G
T
A
C
G
T
A
C
G
T
A
C
G
T
A
C
G
T
A
  
Reverse Opposite:  

A
C
G
T
A
C
G
T
A
C
G
T
A
C
G
T
A
C
G
T
A
C
G
T
A
C
G
T
A
C
G
T
A
C
G
T
A
C
G
T
A
C
G
T
A
C
G
T
  

|  |  |
| --- | --- |
| p-value: | 1e-4 |
| log p-value: | -1.065e+01 |
| Information Content per bp: | 1.530 |
| Number of Target Sequences with motif | 137.0 |
| Percentage of Target Sequences with motif | 1.91% |
| Number of Background Sequences with motif | 1197.7 |
| Percentage of Background Sequences with motif | 1.32% |
| Average Position of motif in Targets | 105.8 +/- 42.1bp |
| Average Position of motif in Background | 101.3 +/- 58.8bp |
| Strand Bias (log2 ratio + to - strand density) | -0.1 |
| Multiplicity (# of sites on avg that occur together) | 1.01 |
| Motif File: | file (matrix) reverse opposite |

#### Similar de novo motifs found

|  |  |  |  |  |  |  |  |
| --- | --- | --- | --- | --- | --- | --- | --- |
| Rank | Match Score | Redundant Motif | P-value | log P-value | % of Targets | % of Background | Motif file |
| 1 | 0.913 | C G T A C G T A C G T A C G T A C G T A C G T A C G T A C G T A C G T A C G T A | 1e-1 | -2.875549 | 57.96% | 57.02% | motif file (matrix) |
| 2 | 0.815 | C G T A C G T A C G T A C G T A G T C A C G T A C G T A C G T A | 1e0 | -1.094314 | 54.39% | 54.14% | motif file (matrix) |
