## Supplementary_Information for "Genome-wide mapping of the *Galleria mellonella* larvae transcription start sites during fungal infection and treatment": motif33.similar.html

### Information for motif33

A
G
T
C
G
T
A
C
A
G
T
C
G
T
C
A
A
G
T
C
A
G
T
C
A
G
T
C
A
G
T
C
A
G
T
C
A
G
T
C
A
G
T
C
A
T
G
C
  
Reverse Opposite:  

A
T
C
G
A
C
T
G
C
T
A
G
A
C
T
G
C
T
A
G
A
C
T
G
A
C
T
G
A
C
T
G
A
C
G
T
A
C
T
G
A
C
T
G
A
C
T
G
  

|  |  |
| --- | --- |
| p-value: | 1e-3 |
| log p-value: | -7.798e+00 |
| Information Content per bp: | 1.890 |
| Number of Target Sequences with motif | 884.0 |
| Percentage of Target Sequences with motif | 12.29% |
| Number of Background Sequences with motif | 10024.8 |
| Percentage of Background Sequences with motif | 11.03% |
| Average Position of motif in Targets | 102.4 +/- 53.6bp |
| Average Position of motif in Background | 99.3 +/- 56.3bp |
| Strand Bias (log2 ratio + to - strand density) | 0.0 |
| Multiplicity (# of sites on avg that occur together) | 1.27 |
| Motif File: | file (matrix) reverse opposite |

#### Similar de novo motifs found

|  |  |  |  |  |  |  |  |
| --- | --- | --- | --- | --- | --- | --- | --- |
| Rank | Match Score | Redundant Motif | P-value | log P-value | % of Targets | % of Background | Motif file |
